# Corticothalamic circuit mechanisms underlying brain region and ageing variations in resting-state alpha activity

**DOI:** 10.1101/2025.08.08.669373

**Authors:** Sorenza P. Bastiaens, Davide Momi, Leanne Rokos, Taha Morshedzadeh, Kevin Kadak, M. Parsa Oveisi, John D. Griffiths

## Abstract

Understanding the neural mechanisms underlying oscillations in resting-state brain activity, which exhibit substantial spatial and age-related variations, remains a significant challenge. This study aims to characterize the contributions of neural circuits to the mechanisms governing resting-state alpha oscillations, which are crucial for various neurocognitive processes and pathologies. Using the Cam-CAN dataset, source-space MEG analyses revealed a pronounced posterior-anterior gradient in alpha frequency, alpha power, and aperiodic components, alongside notable age-related changes. Through neurophysiological modelling, we uncover strong corticothalamic interactions in occipital regions, contrasting with predominantly corticocortical interactions in frontal areas. Ageing is associated with reduced intrathalamic activity and increased corticothalamic delay in occipital regions, while fronto-central regions exhibit increased intrathalamic activity. These findings establish how different circuits shape alpha oscillations across posterior-anterior axis and age, providing a mechanistic foundation for targeted clinical interventions and offering benchmarks for future studies in patient populations.

## 1 Introduction

The processes involved in the emergence of intrinsic brain rhythms have gained significant attention due to their importance in neurocognitive processes [1, 2, 3]. These spontaneously-generated brain rhythms can be captured using magnetoencephalography (MEG) or electroencephalography (EEG), which measure the magnetic and electrical fields produced by synchronized neuronal activity, respectively. The rhythms, categorized into frequency bands associated with specific brain states [4, 5], are characterized by distinct peaks in the M/EEG power spectra. Key power spectral features include periodic activity components, namely peak frequency and power, and the slope of aperiodic activity. During quiet wakefulness, in particular with eyes-closed, the alpha rhythm (8-12Hz) becomes the dominant oscillatory frequency observable in M/EEG data, occurring most strongly in occipital regions. Despite its long-known existence, the mechanisms responsible for alpha generation remain insufficiently understood, and its origin is still debated [6]. Some studies have suggested the thalamus as the primary source of alpha rhythms driven by pulvinar or lateral geniculate nucleus [7, 8, 9]. Other studies emphasized the interaction between the thalamus and the cortex in modulating the alpha rhythm [10, 11, 12]. However, even early studies have shown that normal alpha rhythm can persist even with thalamic damage, suggesting the thalamus is not the only alpha generator, and pointing to a combination of thalamocortical and corticocortical circuit mechanisms [13, 14]. Recent studies have even found that cortical alpha waves lead alpha oscillations in the thalamus [15]. Thus, the alpha rhythms observed in different brain regions may originate from distinct processes, and emerge from multiple cortical and/or thalamic sites. Insights into these differing dynamics are drawn from observing age-related spatial variations in alpha dynamics through power spectral features captured with M/EEG [16, 17].

Previous studies using MEG data have demonstrated spatial variations in resting-state oscillations features, including a significant reduction in the dominant alpha frequency along the posterior-anterior axis. Mahjoory et al. [18] showed that the observed posterior-to-anterior decrease in alpha peak frequency is negatively correlated with cortical thickness, reflecting the brain’s hierarchical organization from early sensory to higher-order associative regions, highlighting a fundamental structure-function relationship in the brain. Furthermore, researchers identified a region-specific arrangement of oscillations characterized by an increase in frequency along both a medial-to-lateral and posterior-to-anterior gradient [19]. Resting-state alpha power maps show the highest values in the occipital cortex, with a gradual decline towards the anterior regions [20]. Studies of resting-state aperiodic activity using MEG and EEG data have identified a pronounced anterior-posterior gradient in the 1/f slope distribution, with steeper slopes in higher frequencies (5-100 Hz and above 15 Hz) in posterior regions and steeper slopes in lower frequencies (0.1-2.5 Hz and below 15 Hz) in the frontal cortex, depending on the specific frequency range examined [21, 22].

Resting-state oscillations not only exhibit spatial variations but also age-related changes. Studies have found that healthy older adults typically show decreased theta and alpha frequencies and powers, alongside increased relative beta and gamma powers, compared to younger individuals [23]. Higher-frequency powers (beta and gamma) mainly increase in rostral regions with age, whereas lower-frequency powers (delta, theta, and alpha) primarily decrease in caudal regions [16]. Analysis of EC resting-state relative power using the Cam-CAN dataset demonstrated a decrease in delta, theta, and alpha relative powers with age [24]. Furthermore, the decomposition of power spectra into periodic and aperiodic components highlighted the need for meticulous spectral component analysis to accurately quantify age-related changes [25]. Research using EEG data showed age-related differences in peak alpha characteristics upon adjusting for aperiodic activity [26]. Older adults displayed reduced peak alpha power and slower peak alpha frequency than younger subjects, although the differences in aperiodic-adjusted peak alpha power were less marked than those observed with unadjusted power. Research also suggests that the aperiodic component (1/f slope) tends to flatten with age, indicating a shift in spectral power distribution towards higher frequencies [27, 28, 29]. It has been suggested that the exponent parameter, representing the negative slope of the power spectrum, reflects shifts in the balance between excitation and inhibition [29, 30], offering significant insights into their observed dynamics.

While alpha frequency and power have been extensively studied across spatial domains, the investigation of aperiodic components in relation to space and age remains underexplored in resting-state EC studies. Similarly, spectral features that change with age are often characterized by discrete spatial points in occipital versus frontal regions, rather than demonstrating a continuous spatial gradient. Previous research has explored alpha frequency and alpha power across different brain regions, such as Park et al. [31]. Their study, however, focused solely on young and old groups and did not consider aperiodic features. Additionally, while Chiang et al. [32] employed a clustering algorithm to assess continuous changes in alpha frequency and power with age, finding that alpha frequency increases until around age 20 before gradually declining, only considering the spatial aspect to a limited extent. A more detailed spatial analysis across brain regions could reveal more intricate patterns of variation, while examining these metrics across distinct age groups would offer a more nuanced understanding of age-related changes. Finally, none of this prior work investigated aperiodic features of the power spectrum in resting-state eyes closed, which is a critical component in the interpretation of inter-subject differences and age-related changes in human electrophysiology datbastia. Thus, one of the aims and novel contributions of the present study is to comprehensively analyze these features in a continuous fashion across the dimensions of age and space (brain region). While these studies shed light on the spatial and age-related changes in alpha activity, they leave open important questions about how these alterations relate to the underlying neurobiological mechanisms that generate and regulate alpha rhythms. To overcome this obstacle, we employ a corticothalamic neural field model developed by Robinson et al. [33], which has been validated extensively and is based on physiological and physical principles [34, 33, 35]. The BrainTrak library [36] provides a fitting algorithm to the model which converts spectral M/EEG experimental data into interpretable parameters that describe neural populations and their connections within the thalamocortical system. This approach builds on the work of van Albada et al. [37], who investigated age-associated trends in physiologically-based EEG spectral parameters at the Cz electrode. Our research extends Van Albada’s previous work by fitting averaged spectra across the entire brain, rather than focusing on a single electrode.

The goal of this study is to characterize circuits involved in generating resting-state oscillations across the brain, in particular alpha rhythms, by identifying distinct spatial and age-related features in MEG data. These features are then correlated with model parameters representing underlying neural dynamics to elucidate key mechanisms, circuits, and parameters involved in alpha rhythmogenesis. More specifically, we aim to assess the roles of cortical and thalamic activity across the brain in shaping alpha rhythms. We analysed the publicly available Cambridge Centre for Ageing Neuroscience (Cam-CAN) [38] resting-state MEG data from 631 subjects, obtaining power spectra for 8196 sources through source localization. Using FOOOF [26], we parameterized the power spectra to separate periodic and aperiodic activity, allowing us to subsequently determine alpha frequency, and power. We also defined aperiodic components for low and high frequencies separately. The cortex was parcellated into 200 regions of interest (ROIs) using the Schaefer atlas [39], and average values for the four features were calculated for each ROI, resulting in 200 values per feature. To examine shifts in corticothalamic model parameter distributions over space, power spectra were averaged and fitted for each of the 200 ROIs. This model has 9 physiological parameters, and also features a reduced form that distils these parameters into three primary circuits (*x* = corticocortical, *y* = corticothalamic, *z* = intrathalamic). We then focused on trends along the posterior-anterior axis for both the empirical features and modelling parameters to evaluate the corticothalamic circuit’s contributions across different spatial regions and age groups. An outline of these analyses is provided in Figure 1, with more detailed information available in the following Methods section.

**Figure 1.**
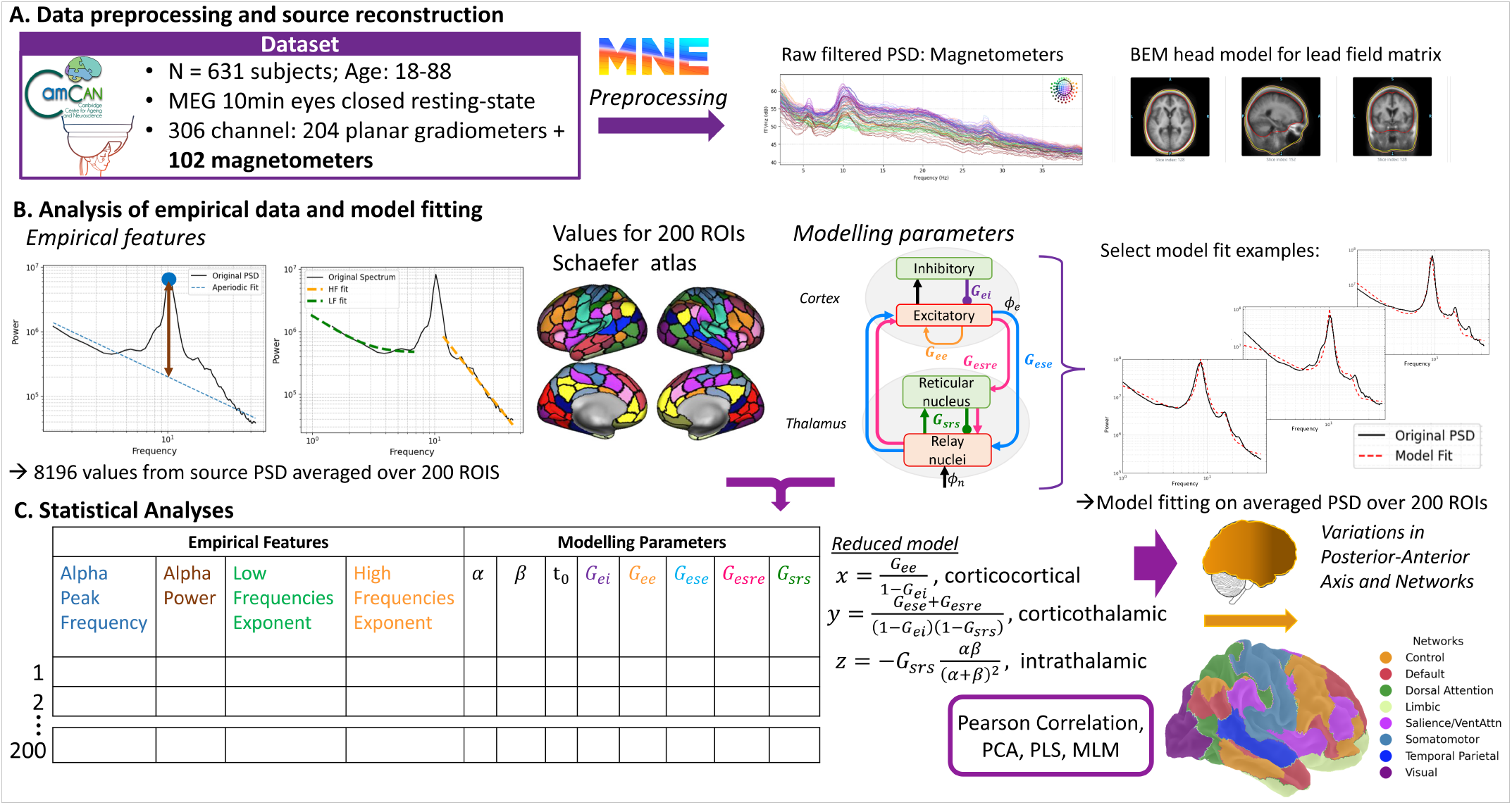
MEG data analysis and physiological modelling methodology. **A)** The Cam-CAN MEG eyes-closed resting-state dataset was preprocessed using the MNE-BIDS pipeline [40], including inverse modelling to obtain source-space power spectra. **B)** From each source power spectra four features of interest (alpha frequency, alpha power, LF aperiodic exponent, HF aperiodic exponent). Alpha frequency and power were obtained using the FOOOF library to separate the periodic and aperiodic activity [26], and HF and LF aperiodic activity were separately estimated with line fitting and monoexponentially decaying function, respectively. The values were then averaged over all dipoles within each of the scale-200 Schaefer atlas parcels. Corticothalamic circuit parameters were estimated by fitting the average power spectra within each parcel using the BrainTrak library [35]. **C)** For the 200 values of the empirical features and modelling parameters, variations across the posterior-anterior axis and age were investigated with Pearson correlation, PCA, and behavioural PLS. Associations between the empirical features and the parameters were studied using MLM.

In summary, we first identify key features in the empirical power spectra from restingstate data, then apply modelling techniques to investigate the role of various corticothalamic circuits. Our findings offer insights into spatial and age-related aperiodic variations, as well as the mechanisms underlying alpha oscillations, by pinpointing the parameters that influence changes in frequency, power, and the aperiodic activity, while highlighting the spatial alterations associated with ageing. We show that the corticothalamic model may analytically capture spatial and ageing variations in resting-state alpha oscillations. By identifying specific markers linked to these variations, we aim to enhance diagnostic tools and develop more targeted clinical treatments for neurological disorders. Furthermore, understanding these patterns is crucial, as it may reveal insights into age-related changes in brain function, guiding the development of strategies to restore normal oscillatory activity and ultimately improving patient outcomes.

## 2 Methods

### 2.1 Subjects and Protocol

The MEG dataset is part of the Cambridge Centre for Ageing and Neuroscience (Cam-CAN) database [41, 38], comprising 9m20s recordings of eyes-closed resting-state data from 631 healthy subjects aged between 18 and 88 years. No exclusion criteria were applied to the data. The data was acquired with a 306 VectorView system (Elekta Neuromag, Helsinki) which measures the magnetic field with 102 magnetometers and 204 orthogonal planar gradiometers. For a more comprehensive understanding of the technical aspects related to MEG instrumentation and data acquisition, we recommend consulting the reference publications pro-vided by the Cam-CAN [38]. The analysis included 607 subjects, due to modelling failures or the absence of an identifiable alpha peak in some subjects.

### 2.2 MEG preprocessing

As this database has been previously investigated, we implemented a publicly available pre-processing protocol proposed by Engemann et al. [42] which uses the MNE-BIDS-Pipeline to perform an automated processing of the MEG data. Management of the pipeline is achieved by using a dataset-specific configuration file outlining the processing steps pertinent to the characteristics of the data for the MNE-BIDS-Pipeline. Notably, the scripts remain unaltered and can be readily applied to a wide range of datasets. The data used was previously maxfiltered with tSSS method that is integrated within the Elekta Neuromag MaxFilter 2.2 software. According to the Cam-CAN data portal, the default settings for MaxFilter 2.2 were applied, encompassing a temporal signal space separation (tSSS) with a correlation threshold of 0.98 and a 10-second window. Bad channel correction was activated, while motion correction remained disabled. A 50Hz+harmonics (mains) notch filter was also applied to mitigate electrical interference. Following the application of tSSS, the signal representation on magnetometers undergoes a linear transformation relative to the signal representation on gradiometers. Consequently, when performing analyses exclusively on either magnetometers or gradiometers, the outcomes are nearly identical, as demonstrated in previous work [43]. To reduce computation times, as in previous analysis, we preprocessed only the magnetometers. The MNE-BIDS pipeline preprocessing steps include bandpass-filtering of raw signals between 0.1 and 49Hz using a zero-phase finite impulse response (FIR) filter with a Hamming window. Source modelling was then performed using a template head model through FreeSurfer. It is worth nothing that, according to [42], the filterbank source model required a reduced-size template head model for the Cam-CAN dataset, created using a scaled MRI (see https://github.com/meeg-ml-benchmarks/brain-age-benchmark-paper). A forward model was computed using a single-compartment boundary element model (BEM), which defines the lead field relating neural sources to the MEG sensor array. For the source space analysis, the inverse operator used was dSPM (default) and the noise covariance was a diagonal ad-hoc. The source-space resolution is of 8196 vertices (oct-6). The output of the pipeline includes the filtered raw data and the inverse operator used to compute the 8196 source PSDs. The source PSDs were computed using the `mne.minimum_norm.compute_source_psd` function with default settings, and the dynamic statistical parametric mapping (dSPM) approach was utilized for source estimation. For power spectral density estimation, a Hanning window was applied to segments of the data, following a method similar to Welch’s technique, which involves averaging the periodograms of the windowed segments.

### 2.3 Empirical Analysis: Estimation of alpha frequency, alpha power and aperiodic activity for low and high frequencies

For each subject and each of the 8196 source power spectra, the following analyses were conducted. First, the power spectra were parametrized using FOOOF v.1.0.0 [26] to separate periodic from aperiodic activity using the ‘fixed’ aperiodic fitting. The alpha frequency peak was then estimated by subtracting the aperiodic fit from the original power spectra, and continuous wavelet transform function from the library SciPy v.1.10.1 [44] was applied to the flattened spectrum for peak detection within the frequency range of interest. This method was chosen over the peak frequency estimation provided by FOOOF for increased precision. The alpha power was directly extracted from the FOOOF object (PW) and corresponded to the power of the most prominent peak within the alpha frequency range, measured above the aperiodic fit. Two measures for aperiodic activity were defined: the exponent high-frequency (HF) aperiodic activity and the exponent for low-frequency (LF) aperiodic activity. Based on data observation, LF aperiodic activity seemed to be best represented by the standard aperiodic FOOOF fitting, whereas HF aperiodic activity was more accurately modelled using the ‘knee’ fit option (Supplementary S.4). However, the knee fitting was not consistently applicable, as some power spectra lacked a distinct knee, and in some LF cases, the higher frequencies skewed the fitting for the lower frequencies. Consequently, in order to estimate the HF aperiodic activity, we applied a linear fit to the log scaled data for frequencies above 12 Hz using the scipy.polyfit function. For the LF aperiodic component, we fitted a monoexponential decaying function to the data up to 7Hz. By fitting the exponent of the aperiodic activity separately for low and high frequencies, we aimed to capture the specificity of the slopes for each frequency range more accurately. This resulted in four different features characterizing the power spectra for empirical analysis. Finally, the cortex was parceled into 200 regions-of-interest (ROI) based on the Schaefer atlas with 17 networks [39]. The values of the four features for each ROI were determined by averaging the estimated values across the vertices within the corresponding region. To investigate changes along the posterior-anterior gradient, the y-coordinate of the MNI space for each ROI was extracted using the source estimate file, resulting in 200 values per subject for each feature for comparison with the modelling parameters.

### 2.4 Modelling: Corticothalamic neural population model

In order to investigate the involvement of cortical or corticothalamic circuits in resting-state oscillations across the cortex, we opted for a well-established corticothalamic neural field model [34, 45, 33, 46, 47, 48, 49, 36]. The model is composed of four neural populations, two cortical populations (excitatory *e* and inhibitory *i* cortical neurons), and two thalamic populations (inhibitory thalamic reticular nucleus *r*, and excitatory thalamic relay *s*). In this model, cell firing acts as sources of pulse fields, representing the average spike rate in each population. These spike rate fields are then filtered by dendrites in postsynaptic neurons before reaching the cell body. Subsequently, the membrane potential influences the population’s firing rate. Therefore, the aggregate activity of each neural population (*a* = *e, i, r* or *s*) is described in terms of firing rate (*Q*_*a*_), and membrane potential (*V*_*a*_). *V*_*a*_ is transformed into *Q*_*a*_ by a nonlinear sigmoid function defined as

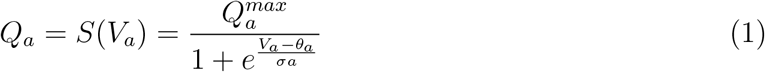

with *θ* as the mean threshold voltage,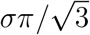 the standard deviation of the threshold distribution, and *Q*_*max*_ the maximum firing rate. *V*_*a*_ is determined by the contributions of *ϕ*_*b*_ (which are the mean density of outgoing spikes produced by population b) from presynaptic populations, the strength of the connections (*ν*_*ab*_) and the smoothing effects resulting from synaptodendritic dynamics and soma capacitance (defined by *D*_*α*_). This results in the following equations:

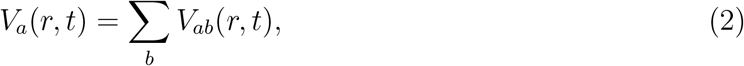

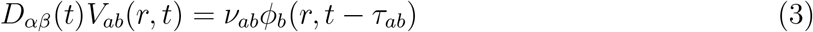

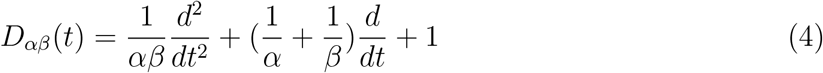

Here, *ν*_*ab*_, the connection strength, is equal to *s*_*ab*_*N*_*ab*_ where *N*_*ab*_ represents the average number of incoming synapses per neuron, *s*_*ab*_ is the average time-integrated strength of the soma response per incoming spike rate [49, 36]. All connections are excitatory (positive) except for *ν*_*ii*_, *ν*_*ei*_ and *ν*_*sr*_ which are inhibitory (negative). The parameters 1*/β* and 1*/α* respectively represent the characteristic rise time and decay time of the response at the cell body. Although these parameters can vary for each connection in the model, a weighted average over the corticothalamic system is used to provide a single effective value while preserving the dynamics. Time delays *τ*_*ab*_ are due to discrete anatomical separations between populations. The only nonzero time delays are connections between cortical populations and thalamic populations which are equal to *t*_0_*/*2. This corresponds to the corticothalamic and thalamocortical propagation time. The outgoing action potentials propagating to the next neural population, represented as a field *ϕ*_*a*_(*r, t*), can be approximated by a damped wave equation with a source term *Q*_*a*_ [34, 33] *where*

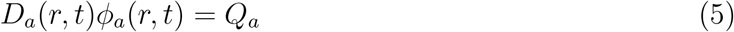

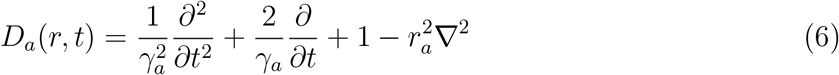

with *r*_*a*_ as the mean range, *γ*_*a*_ as *v*_*a*_*/r*_*a*_ is the temporal damping rate defined by the propagation velocity (*v*_*a*_). In this model, only *r*_*e*_ is considered large enough to produce significant propagation effects. For the other populations (*a* = *i, r, s*), owing to the short range of cortical inhibitory axons and the relative smallness of the thalamus, the approximation *ϕ*_*a*_ = *Q*_*a*_ = *S*[*V*_*a*_] can be made. The model can be further simplified by assuming random corticocortical connectivity, where the number of connections between populations is proportional to the number of synapses. This implies that the connection strengths are symmetric, resulting in a model composed of eight independent connections [36]. The EEG power spectrum is defined by considering the spatially uniform steady states of the model, and taking the Fourier transform of equation 3, with the firing rate *ϕ*_*e*_ expressed as

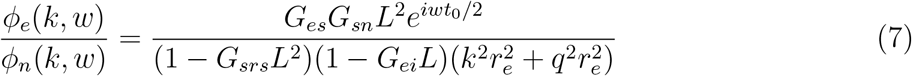

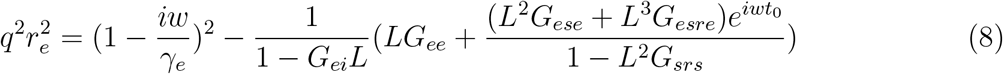

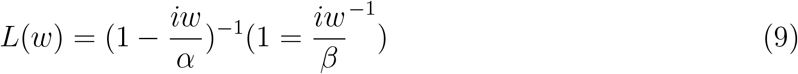

as described in Robinson et al. [33] where *k* and *w* are the wave vector (with magnitude *k* = 2*π/λ* where *λ* is the wavelength) and angular frequency (*w* = 2*πf* where *f* is the cyclic frequency in Hz), respectively. *ϕ*_*n*_ corresponds to the external signal coming into the thalamic relay and is taken to be white noise. With uniform spectral power density for nonzero *k* and *w*, in the linear approximation the value of *ϕ*_*n*_(*k, w*) influences only the normalization of the power spectrum. Since the steady-state firing rates depend solely on the product 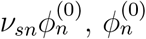 is set to 1*s*^*−*1^ without loss of generality when calculating the steady-state firing rates. *G*_*ab*_ = *ρ*_*a*_*ν*_*ab*_ is the gain which represents the response in neurons *a* due to unit input from neurons 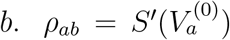 and is obtained by linear approximation. The overall gain quantities for the excitatory corticothalamic, inhibitory corticothalamic, and intrathalamic loops are *G*_*ese*_ = *G*_*es*_*G*_*se*_, *G*_*esre*_ = *G*_*es*_*G*_*sr*_*G*_*re*_, and *G*_*srs*_ = *G*_*sr*_*G*_*rs*_, respectively.

The power spectrum is then obtained by integrating *ϕ*_*e*_(*k, w*) over k. In order to account for the finite size of the brain, the cortex is modelled as a rectangular sheet. Periodic boundary conditions are applied to simplify the numerical investigation of the modal structure of the power spectrum, though this choice is not crucial to the system’s dynamics [33]. Under periodic boundary conditions, the power spectrum is expressed as:

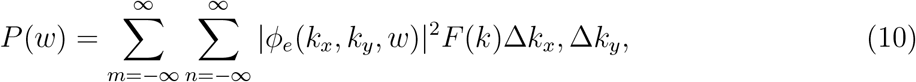

with

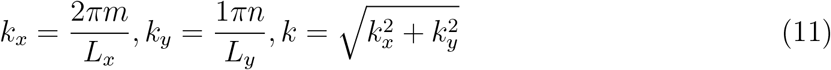

The dimensions of the two-dimensional rectangular cortex are *L*_*x*_ ⨯ *L*_*y*_ with *L*_*x*_ = *L*_*y*_ = 0.5*m* [33]. The filter function *F* (*k*) approximates the low pass spatial filtering effects due to volume conduction by the cerebrospinal fluid, skull, and scalp. It is defined as:

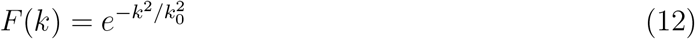

with a low-pass cutoff at *k*_0_ ≈10*m*^*−*1^ based on the spherical harmonic head transfer function developed by Srinivasan et al. [50].

The PSD depends on the five gains *G*_*ee*_, *G*_*ei*_, *G*_*ese*_, *G*_*esre*_, and *G*_*srs*_. In the stable regions of the parameter space at low frequencies, a reduced 3-dimensional space can be defined to represent the model parameters. In this reduced space we have:

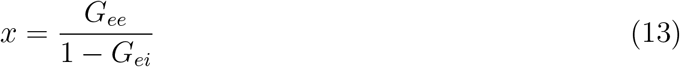

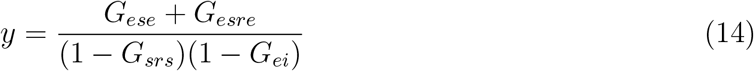

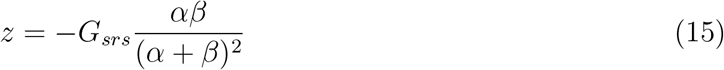

which parametrize cortical, corticothalamic, and intrathalamic loop gains, respectively. These capture the most important dynamics of the model. From these equations, the reduced model based completely on the xyz quantities can be derived. The details are presented in Abeysuriya et al. [35].

To reduce the impact of high-frequency EMG artifacts from pericranial, cervical, and extraocular muscles on the higher frequencies of the power spectra, the amplitude of the electromyographic artifact correction *A*_*EMG*_ is incorporated into the PSD. This is done according to the following equation:

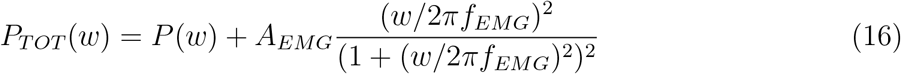

where the term *A*_*EMG*_ is fitted along with the other parameters for *f*_*EMG*_ in range of 10 to 50Hz.

The BrainTrak corticothalamic neural field parameter estimation toolbox in MATLAB was used to fit the empirical power spectra. With this toolbox, the linearized corticothalamic model is fitted, yielding gain parameters, *α, β, t*_0_ and *A*_*EMG*_. The reduced model’s parameters *x, y* and *z*, representing the cortico-cortical, corticothalamic and intrathalamic loops, respectively, are then calculated from the fitting. The source power spectra are averaged for each ROI, resulting in 200 power spectra per subject, which are then fitted yielding 200 values for each parameter. Similarly to the empirical analysis, we then have a value for each ROI. The BrainTrak toolbox uses the Metropolis-Hastings algorithm, a Markov Chain Monte Carlo (MCMC) method for model fitting by providing samples from general probability distributions. The parameters were constrained within the stability limits established in previous literature [36] to ensure the biological plausibility of the fitted parameters. Additionally, the gain parameters for all connections were capped at 20 (|*G*_*ab*_|*<* 20) to minimize the model’s sensitivity to input noise. The chi-squared *χ*^2^ error for model optimization is used, calculated between the 1 and 45Hz frequency bins. The aim is to reduce the error between the empirical power spectra and the simulated model power spectra. The parameter space is initially explored through a random walk of 100,000 steps with large step sizes. This process identifies the top points that bring us close to the target values. Following this ‘burn-in’ period, BrainTrak takes smaller steps to more accurately approximate the ground truth.

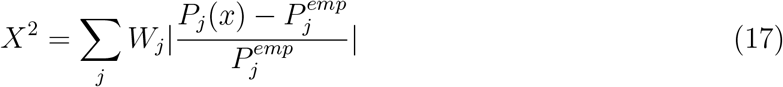

where *P*_*j*_(*x*) is the model prediction and 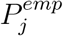 is the empirical data, and j indexes frequency bins from the fast Fourier transform of the empirical data [35]. The term *W*_*j*_ acts as a scaling factor to amplify the influence of lower frequency bands relative to higher frequency bands (*W*_*j*_ *∝ f*^*−*1^). This adjustment enhances the optimizer’s sensitivity to high frequency bins in the power spectrum while minimizing its sensitivity to lower frequencies. This approach is beneficial for reducing artifacts in MEG data, as the primary artifacts affecting the 1-45 Hz range, such as movement, and eye blink artifacts, produce low-frequency noise that needs to be mitigated.

### 2.5 MEG power spectra dynamics are robustly captured by aperiodic and corticothalamic model fitting across subjects

We report here the R-squared values for both the aperiodic and corticothalamic model fittings, highlighting the overall goodness-of-fit across subjects. As mentioned previously, in order to describe the aperiodic activity, we separately estimated HF and LF slope as the spectra showed a reliable change in slope between 1-7Hz (before alpha peak), and above 12Hz (after alpha peak). For the HF range, a linear regression using a polynomial fit was applied to 12Hz and above in the log scale, yielding a mean R-squared of 0.917 (SD = 0.092) across subjects, indicative of a consistently good fit. The 95% confidence interval for this fit was narrow, from 0.9174 to 0.9175, underscoring its robustness. In the 1-7Hz range, we applied a monoexponential decay function to capture the aperiodic activity of lower frequencies, which yielded a slightly lower mean R-squared of 0.891 with a standard deviation of 0.172, reflecting greater variability across subjects. The 95% confidence interval ranged from 0.8913 to 0.8916. Notably, fitting errors occurred in 68 subjects for some of the power spectra within the LF range, due to deviations from the expected exponential decay, such as the presence of a bump or a more linear trend at lower frequencies. These were dropped from the parcel average.

The corticothalamic model fitting of the power spectra was successfully achieved across 609 subjects. To assess the overall fit, we calculated the mean likelihood value for each subject and then averaged across all subjects. The mean likelihood estimate across subjects was 0.80 (SD = 0.063) indicating a generally good fit of power spectra for each subject.

### 2.6 Statistical Analyses

We initially explored the correlation between empirical features and modelling results along the spatial posterior-anterior axis. Individual Pearson correlations were estimated for all empirical features and modelling parameters. Additionally, we conducted a population-level analysis by dividing the subjects into seven age groups (18-27, 28-37, 38-47, 48-57, 58-67, 68-77, 78-88). For each region of interest (ROI), we averaged the values across subjects within each age group. We then carried out separate principal component analyses (PCA) on the empirical features and fitted model parameters to identify the principal components and their loadings across various brain regions. behavioural partial least squares (PLS) correlations, a multivariate statistical method, were performed to analyse the relationship between a predictor variable (in this case, age) and features (e.g., neural activity metrics) across different ROIs [51, 52] on MATLAB. PLS analysis allows us to extract latent variables that maximize the covariance between age and the features, providing insights into how empirical features relates with age across different brain regions. Bootstrapping was used to test the reliability of the results, and the significance of the latent variable was determined by permutation testing providing a p-value. Combining the two, we can identify the regions reliably and significantly showing the relationship. We conducted separate PLS analyses for each empirical feature and parameter, as well as combined PLS analyses for all empirical features and for all modelling parameters. We present in the main text the results from the corticothalamic reduced model, but have provided the analysis from the non-reduced model in Supplementary S.2. Mixed linear modelling was used to relate empirical features to modelling parameters taking age and space as fixed effects with the library statsmodels v0.14.0. We built a model for each empirical feature of interest, using it as the response variable, and investigated the interaction between age and region with the parameters of the reduced model. In other words, how the effect of *x, y, z*, and *t*_0_ varies with age and with space. Each participant is assumed to have a random effect on the intercept of the model. This accounts for the fact that measurements within the same participant may be more similar to each other than to measurements from different participants. The equation is as follows:

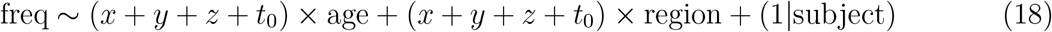

The coefficients for the main effects (modelling parameters) and interaction terms age and space indicate how changes in these modelling parameters and their interactions with age and region are associated with changes in the empirical feature. For example, a positive coefficient for *x* : *age* indicates that as age increases, the effect of *x* on the response variable also increases. In summary, the mixed linear model is used to understand how the modelling parameters and their interactions with age and space explain the variation in the corresponding response variable (which is one of the empirical feature), while accounting for random variability between participants.

## 3 Results

### 3.1 Posterior-anterior spatial gradients are observed in spectral parametrization features

The analysis of four empirical spectral features revealed consistent posterior–anterior gradients in younger adults, which become progressively altered with age. This age-related change reduces the overall monotonic spatial organization, driven by divergent trends between occipital and fronto-central regions. Alpha power exhibited a strong posterior–anterior gradient that remained largely stable across age. In contrast, alpha frequency showed the most pronounced age-related change, with significant decreases in occipital regions. Aperiodic activity was quantified by separating the spectral slope into two components: the high-frequency (HF) slope, defined as the linear fit of the spectrum above the alpha peak (greater than 12 Hz), and the low-frequency (LF) slope, estimated within the 1–7 Hz range. The HF aperiodic slope displayed opposing spatial trends, declining in posterior areas while increasing fronto-centrally with age, resulting in greater spatial dispersion.

A significant Pearson correlation with the MNI y-coordinate, corresponding to the posterior-anterior axis, was observed in 74.14% of subjects for alpha frequency, 98.02% for alpha power, 92.92% for the HF aperiodic slope, and 80.23% for the LF aperiodic exponent (all *p <* 0.005). Averaging feature values within each age decile (18-27, 28-37,…, 78-88), shown in Figure 2A, visually reveals posterior-anterior gradient. To take into account multiple comparisons, the padjusted value is approximately 0.0018 with Bonferroni correction considering 28 comparisons (4 features for 7 age groups). Significant negative correlations were observed in all age groups between MNI y and alpha frequency (*r* = −0.888 to *r* = −0.535, *p <<* 0.001), alpha power (*r* = −0.954 to *r* = −0.924, *p <<* 0.001), HF aperiodic exponent (*r* = −0.859 to *r* = −0.589, *p <<* 0.001), and LF exponent (*r* = −0.862 to *r* = −0.496, *p <<* 0.001) implying that these features decrease from the back to the front of the head. Notably, the correlation strength with the posterior-anterior coordinate was weaker in older subjects for the HF aperiodic exponent and stronger for the LF aperiodic exponent. These consistent spatial variations were observed for all spectral features, with the strongest effects seen in alpha power.

**Figure 2.**
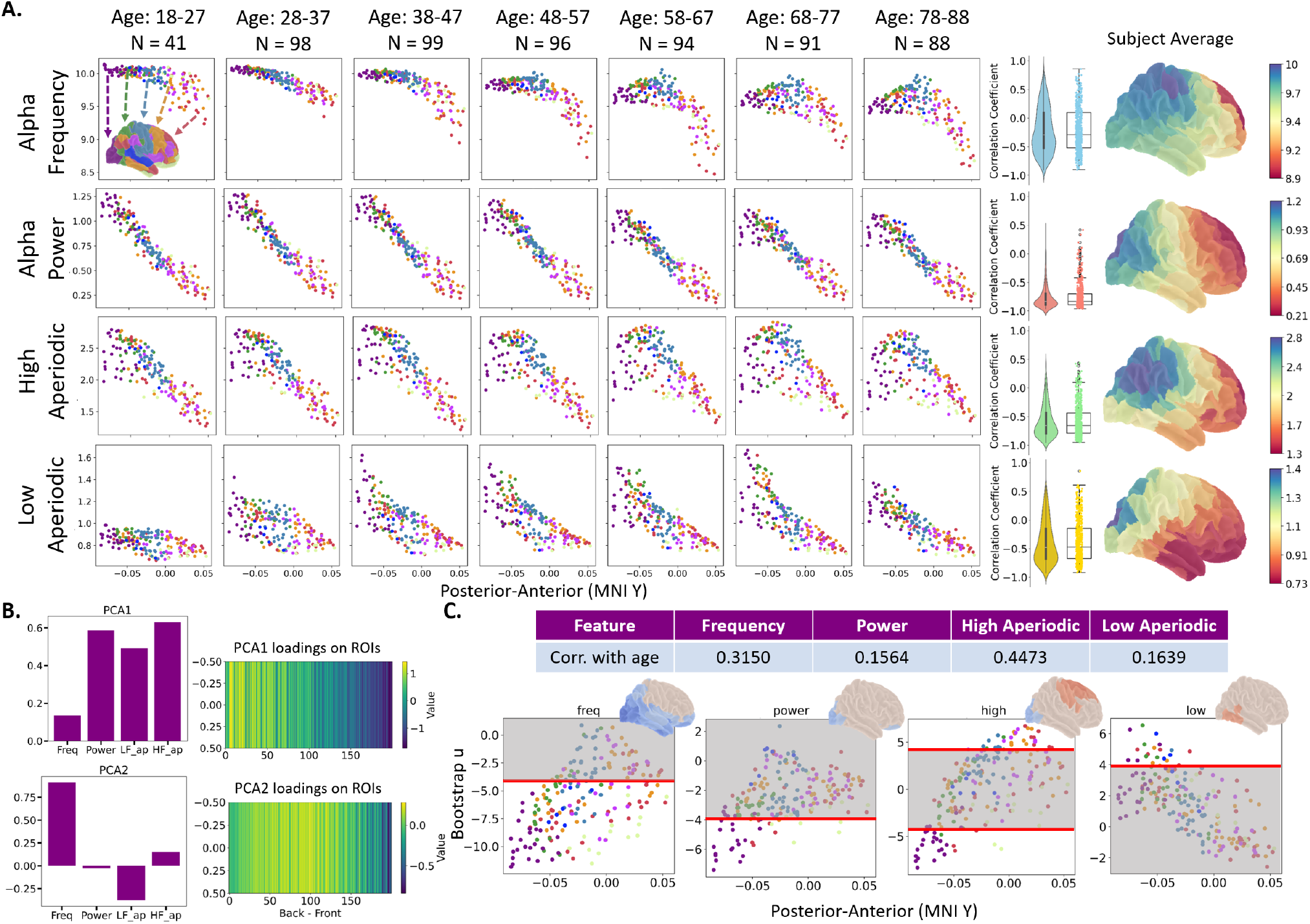
Spatial gradient of empirical features along the posterior-anterior axis with ageing variations. **A)** Empirical features as a function of posterior-anterior axis (MNI y coordinate) across 7 age groups. A strong linear relationship is observed with alpha power across the age groups. A similar trend is observed with HF aperiodic with some spreading occurring with ageing. The age-related decrease in alpha frequency is most pronounced in occipital regions and is accompanied by a slight increase in the low-frequency aperiodic component. **B)** PCA analysis loadings for the first two components and loadings across space. First component reveals the spatial gradient seen with alpha power, HF aperiodic, and LF aperiodic which suggest these features are correlated. Second component shows the trend of frequency across space mostly seen in posterior and central regions. **C)** Correlations identified by the latent variable with behavioural PLS analyses with age as predictor and associated bootstrap ratios. Ratios above 4 show a reliable positive association with age; ratios below −4 show a reliable negative association. Otherwise, the association is not reliably seen with age. Frequency, followed by alpha power, have the most regions reliably showing a negative association with age. Occipital regions show a negative association with age for the HF aperiodic component whereas central-frontal regions show a positive association with age.

To further assess relationships between empirical features and spatial dimensions, we performed PCA to examine the loadings of empirical features and evaluate the strength of the identified components across spatial regions, as shown in Figure 2B. The analysis revealed that the first two principal components accounted for 74% of the total variance in the dataset. The first component was heavily loaded on alpha power (0.59), HF aperiodic exponent (0.49), and LF aperiodic exponent (0.63), suggesting a relationship between alpha power and the aperiodic activity. The second component was heavily positively loaded on alpha frequency (0.90) and negatively loaded on LF exponent (−0.38). Observing the loadings of the first component over space revealed a gradual shift from high to low loadings of the first component, confirming the trends observed previously: Alpha power and aperiodic exponents were more pronounced in caudal regions compared to rostral regions. Spatially, the second component (positively loaded on frequency) exhibited high loadings in occipital-central regions and low loadings in frontal regions, indicating higher frequencies in caudal regions and lower alpha frequencies in rostral regions. The PCA analysis described a pattern implicating power and aperiodic components together, suggesting that these parameters are coupled and exhibit a gradual shift from occipital to frontal regions. Although the frequency shift was slightly less linear in space, it still showed a decreasing trend (Table 1 and Figure 2A). Additionally, an inverse association was identified between alpha frequency and the LF aperiodic exponent.

**Table 1.** Distribution of participants from Cam-CAN dataset across age and sex. Bracketed values reflect the subset retained for analysis after excluding individuals due to failed model fitting or absence of an identifiable alpha peak.

| Age Range | Number of Males | Number of Females | Total |
| --- | --- | --- | --- |
| 18 - 27 | 23 (22) | 24 (19) | 47 (41) |
| 28 - 37 | 50 (49) | 50 (49) | 100 (98) |
| 38 - 47 | 51 (49) | 51 (50) | 102 (99) |
| 48 - 57 | 51 (50) | 48 (46) | 99 (96) |
| 58 - 67 | 47 (46) | 49 (48) | 96 (94) |
| 68 - 77 | 49 (47) | 47 (44) | 96 (91) |
| 78 - 88 | 49 (46) | 42 (42) | 91 (88) |
| Total | 320 (309) | 310 (298) | 631 (607) |

**Table 2.** Summary of Empirical Population-level Pearson Correlation Results.

| Age Range | Alpha PF | Alpha Power | Aperiodic HF | Aperiodic LF |
| --- | --- | --- | --- | --- |
| 18 - 27 | $r = -0.702$ | $r = -0.954$ | $r = -0.855$ | $r = -0.496$ |
| 28 - 37 | $r = -0.888$ | $r = -0.950$ | $r = -0.856$ | $r = -0.588$ |
| 38 - 47 | $r = -0.791$ | $r = -0.950$ | $r = -0.859$ | $r = -0.726$ |
| 48 - 57 | $r = -0.736$ | $r = -0.942$ | $r = -0.792$ | $r = -0.783$ |
| 58 - 67 | $r = -0.694$ | $r = -0.941$ | $r = -0.684$ | $r = -0.791$ |
| 68 - 77 | $r = -0.534$ | $r = -0.924$ | $r = -0.593$ | $r = -0.809$ |
| 78 - 88 | $r = -0.594$ | $r = -0.929$ | $r = -0.589$ | $r = -0.862$ |
| p-value | $p < < 0.001$ | $p < < 0.001$ | $p < < 0.001$ | $p < < 0.001$ |

**Table 3.**
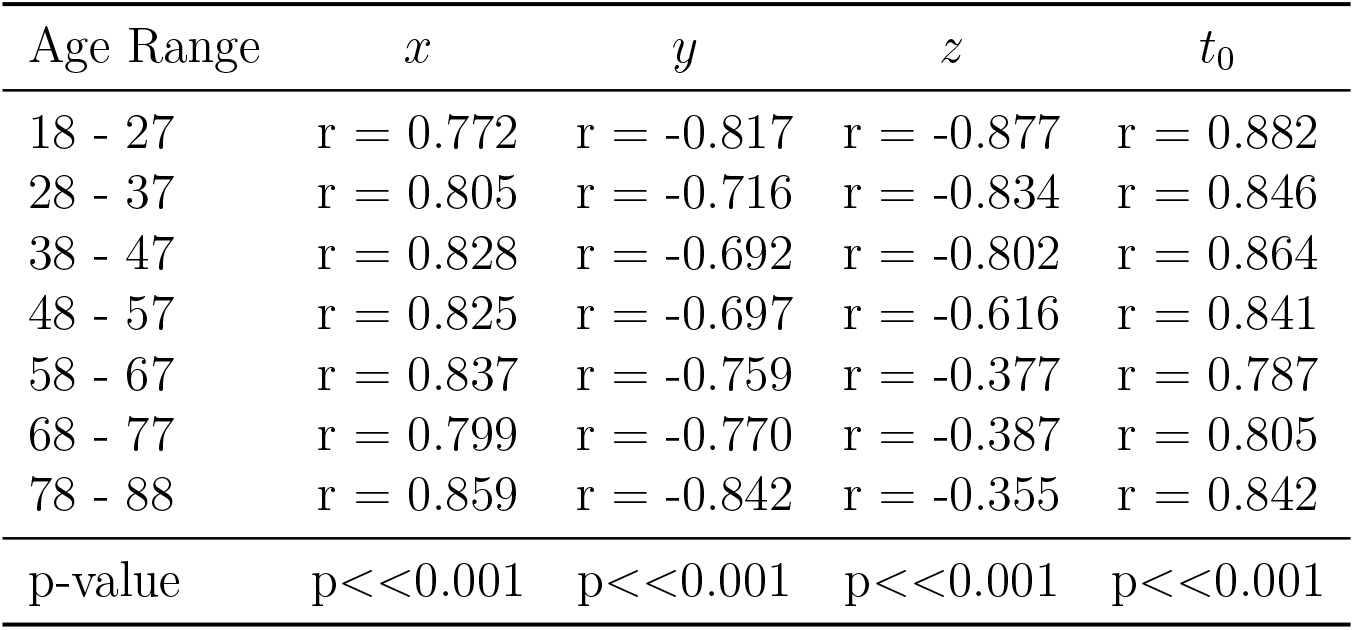
Summary of Modelling Population-level Pearson Correlation Results.

### 3.2 Ageing drives posterior declines in oscillatory activity and anterior increases in aperiodic activity

We conducted behavioural partial least squares (PLS) analyses to explore how each empirical feature relates to age, with spatial variation across the cortex taken into account. The bootstrap ratios of the PLS analyses are presented in Figure 2C. Alpha frequency exhibited age-related changes in the greatest number of brain regions, with 131 regions reliably showing a relationship with age. This was followed by the HF aperiodic slope in 65 regions, then alpha power in 28 regions, and the LF aperiodic slope in 20 regions. For all of them, permutation testing revealed the identification of significant latent variables (*p <<* 0.001), except for LF aperiodic exponent (*p* = 0.0110) suggesting that the association between LF aperiodic slope and age is less pronounced but still present. A stronger, reliable negative relationship between alpha frequency and ageing was observed in occipital regions, particularly within the visual network, indicating that frequency decreases with age. Similar patterns were seen for the HF aperiodic component. The limbic system also showed a decrease in both frequency and power with age. Additionally, a few central-frontal regions demonstrated a positive association between age and the HF aperiodic component. Furthermore, a behavioural PLS analysis of the aggregated empirical metrics revealed two significant latent variables (*p <<* 0.001; Figure 5A). The first latent variable is highly positively loaded with frequency followed by power and a negative loading of LF aperiodic. The second latent variable is positively loaded with HF aperiodic while all other components are negatively loaded. Similar patterns in bootstrap ratios emerge between the individual empirical feature PLS vs. multiple input PLS. Namely, we can say that the first latent variable present the association with age for caudal regions with marked decrease in frequency and power, whereas the second latent variable present the association for central-frontal regions with increased HF aperiodic exponent, identifying distinct spatial empirical changes with age.

### 3.3 Posterior-anterior spatial gradients are also observed in corticothalamic model parameters

Following the analysis of empirical spectral features, we applied the same statistical approach to the model-derived parameters, with the model fit across 200 cortical ROIs. Below, we report the results for the reduced parameter set representing the three main circuits of the model (corticocortical = *x*, corticothalamic = *y*, intrathalamic = *z*) and the corticothalamic delay (*t*_0_). Results from the full parameter model are provided in Supplementary Section S.1. As with the empirical features, model-derived parameters revealed posterior–anterior gradients that varied with age (Figure 3A). Corticocortical and corticothalamic activity followed opposing spatial profiles, with elevated corticothalamic activity in occipital regions relative to frontal areas.These patterns remained largely unchanged across the age groups, suggesting they reflect stable regional specializations in circuit dynamics. In contrast, intrathalamic activity increased with age and displayed a broader, less linear spatial distribution. Finally, the corticothalamic delay decreased in occipital areas with ageing.

**Figure 3.**
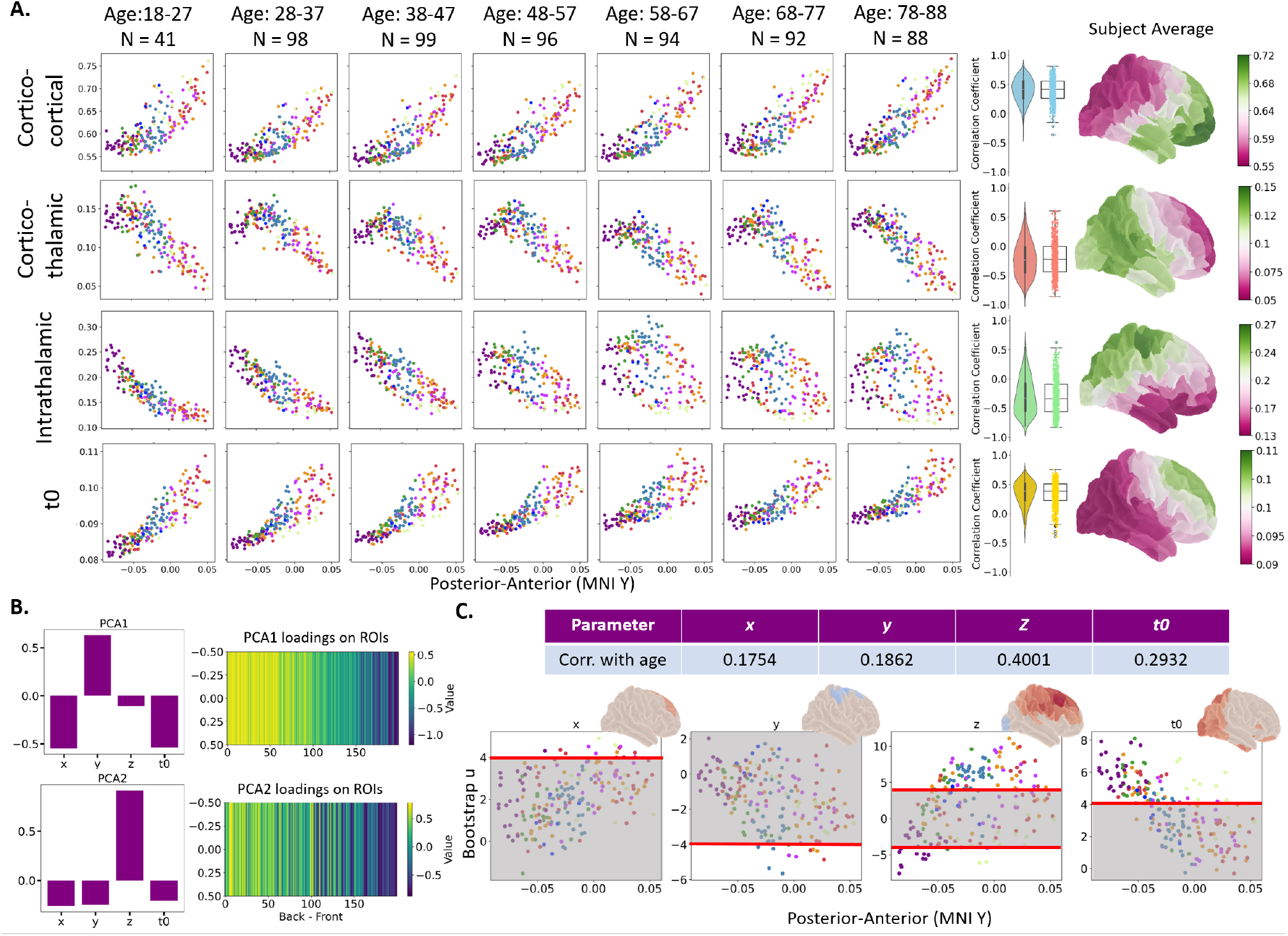
Spatial gradient of modelling parameters along the posterior-anterior axis with ageing variations. **A)** Corticothalamic model parameters as a function of posterior-anterior axis (MNI y coordinate) across 7 age groups. Corticocortical parameter *x* is higher in central-frontal regions, whereas corticothalamic *y* tends to be higher in occipital regions. Intrathalamic parameter *z* presents a strong negative linear relationship that is reduced with age. Corticothalamic delay *t*_0_ decreases mostly in occipital regions with age. **B)** PCA components and loadings over space as a function of the MNI coordinate y (posterior-anterior gradient). The first component presents an anticorrelation between *x* and *y* which are negatively and positively loaded respectively. The component has a spatial gradient with high values in occipital regions and lowest in central-frontal regions. The second component is mainly positively loaded with *z* that is more prominent in occipital-central regions. **C)** Correlations identified by the latent variable with behavioural PLS analyses with age as predictor and associated bootstrap ratios.. Ratios above 4 show a reliable positive association with age; ratios below −4 show a reliable negative association. Otherwise, the association is not reliably seen with age. *t*_0_ presents a positive association with age mostly in occipital regions and in the limbic system. *z* presents a positive association with age for central-frontal regions.

More specifically, pearson correlation analyses revealed significant correlations between spatial coordinates and the reduced model parameter *x* in 84.51% of the subjects, *y* in 64.09%, and *z* in 71.00%. The results of the full model showed the highest correlation for *α* in 95.06%, followed by *G*_*srs*_ in 81.55%, then *t*_0_ in 79.41%, and *G*_*ese*_ in 72.65%. All other parameters showed a spatial gradient in less than 50% of the subjects, except for EMG, which is consistent with the presence of additional artifacts in frontal regions. For further details on these parameters please refer to S.1. By separating in different age group, we observed opposite trends for parameters *x* and *y* in space with their correlation coefficients remaining relatively stable as presented in Figure 3A. Furthermore, higher inhibitory intrathalamic *z* and excitatory corticothalamic activity *y* are present in occipital regions (Figure 3A). However, parameter *z* showed a significant decline in the correlation coefficients, with *r* = −0.877 for the younger population to *r* = 0.355 in the older group, indicating a loss of correlation with age. These correlations indicate a posterior–anterior gradient, characterized by stronger corticothalamic and intrathalamic activities in occipital regions and greater corticocortical activity in frontal regions, with intrathalamic activity exhibiting significant age-related changes. We also noticed that *t*_0_ generally increases with age, particularly in occipital regions where the change is more pronounced. PCA of parameters *x, y, z* and *t*_0_ showed that the first two components explain 68% of the variance. The first component exhibited positive loadings for *x* and *t*_0_ but negative loadings for *y*, suggesting a pattern in which these two are anticorrelated (Figure 3B). Spatially, this component showed a smooth posterior-anterior gradient, with higher *y* loadings in occipital regions, and higher *x* and *t*_0_ loadings in frontal regions. The second component had a strong positive loading in *z*, with slight negative loadings in the other three parameters. Although the spatial gradient presented a less smooth transition than the first component, occipital regions generally had higher loadings of *z*. These findings are consistent with the Pearson correlation analysis, which identified *x* and *y* as stable markers across space. Additionally, PCA including all gains and other parameters, as detailed in the S.2, revealed a spatial pattern involving *G*_*srs*_ and *α*.

### 3.4 Intrathalamic gain and corticothalamic delays show distinct regional associations with age

After characterizing spatial gradients, we assessed how these model parameters change with age across cortical regions. The behavioural PLS analysis on the model parameters did not identify consistent association with age for *x* (*p <<* 0.001) and *y* (*p* = 0.002). Only 13 and 9 regions, respectively, showed a significant and reliable relationship, as indicated by the bootstrap ratios (Figure 3C). In contrast, parameters *z* and *t*_0_ reliably showed a relationship with age for 103 and 89 regions, respectively. Parameter *z* exhibited a negative association with age in visual network regions and a positive association in all other regions (*p <<* 0.001). Parameter *t*_0_ showed a positive relationship with age, particularly evident in occipital regions, and a similar trend was observed in the limbic system (*p <<* 0.001).

A similar pattern emerged when the four modelling parameters were combined in a single PLS analysis (Figure 5B), revealing two significant latent variables (*p <<* 0.001). The first latent variable was predominantly negatively loaded with *z* and *t*_0_, while the second latent variable showed a negative loading in *z*, and a positive loading in *t*_0_. This implies that these two parameters are more closely associated with age. The first latent variable showed significant changes in central-frontal regions, whereas the second latent variable was associated with occipital regions and the limbic system.

### 3.5 Alpha power and aperiodic slopes are modulated by corticothalamic and intrathalamic gains, respectively

Associations between empirical features and modelling parameters were explored, first by computing the Pearson correlation coefficient, *r*, between each pair across all brain sources, as shown in Figure 4. Among the gains and empirical features, the strongest positive correlations were observed between *G*_*srs*_ and *α* (*r* = 0.66), followed by alpha power and *G*_*ese*_ (*r* = 0.61), and finally alpha power and HF aperiodic exponent (*r* = 0.59). The strongest negative correlations were between *G*_*ei*_ and *G*_*ee*_ (*r* = *−*0.88), followed by alpha power and *α* (*r* = *−*0.68), then *G*_*srs*_ and HF aperiodic (*r* = *−*0.60), and *t*_0_ and alpha power (*r* = *−*0.47). For the reduced model parameters, *x* was negatively correlated with alpha power (*r* = *−*0.53) and HF aperiodic (*r* = *−*0.49). *y* was positively related to alpha power (*r* = 0.54), suggesting that alpha power drives the anticorrelation between *x* and *y* across space. *z* showed a positive association with HF aperiodic (*r* = 0.45). Alpha frequency showed comparatively weaker correlations, with the highest associations observed for *z* (*r* = 0.23) and *t*_0_ (*r* = *−*0.19). This reduced correlation likely reflects the less linear, more quadratic spatial distribution of alpha frequency, which may limit the ability of PC to fully capture its relationship with the model parameters.

**Figure 4.**
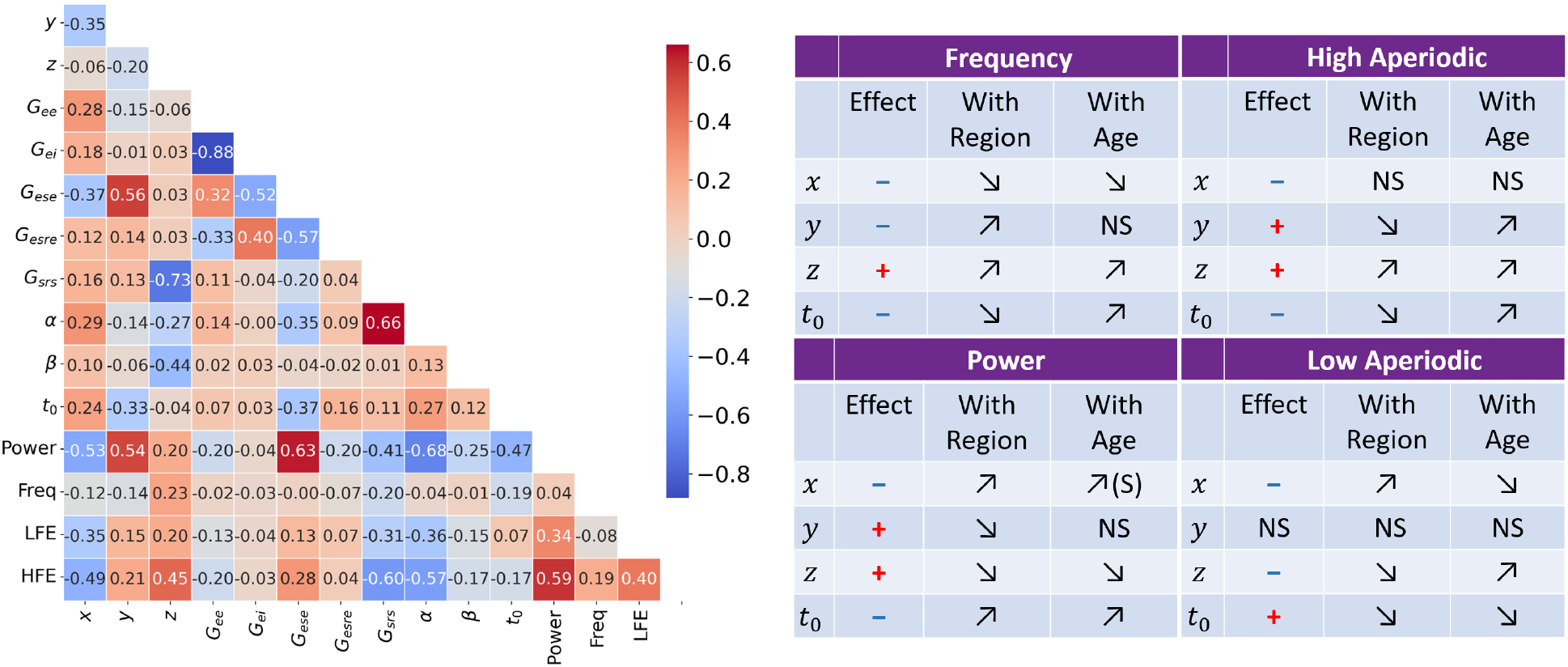
Relationship between empirical features and modelling parameters. **A)** Pearson correlation coefficients with p-value between empirical features and modelling parameters. Alpha power is correlated with *y* (*r* = 0.54), *G*_*ese*_ (*r* = 0.63), and anticorrelated with *x* (*r* = *−*0.53), *α* (*r* = *−*0.68) and *t*_0_ (*r* = *−*0.47). HF aperiodic is correlated with *z* (*r* = 0.45, and thus anticorrelated with *G*_*srs*_ and *α*), and anticorrelated with *x*. Parameters have lower correlation with alpha frequency and LF aperiodic which might indicate that they are not correlated in every brain regions. Strongest correlation is seen with parameters and empirical features that also have a strong spatial correlation. **B)** Summary of the MLM results with age and space as fixed effects investigating the effect of the model parameters on each empirical features. The sign associated with a parameter indicates whether the effect is positive or negative, and the arrows define whether the effect strengthens (up) or weakens (down) with age or across regions (going from back to front). Parameters *z* and *t*_0_ have increased effects on frequency and HF aperiodic with age, highlighting their effect on these components on the ageing variations. *x* and *y* have anticorrelated effect in space on alpha power and HF aperiodic.

### 3.6 Age effects in circuit parameters and spectral features are explained by two distinct latent processes

To further investigate how the association between modelling parameters and empirical features varies by space and age, we performed separate MLM analyses for each empirical feature, treating age and region as fixed effects. Our analysis focused on the parameters of the reduced model and *t*_0_. The effects of the modelling parameters on the empirical features are summarized in Figure 4, and details of the results are provided in Supplementary S.3. For each empirical feature the strength of the effect of at least one of the two parameters *z* or *t*_0_ increased with age. MLM of alpha power and the HF aperiodic exponent showed a negative correlation with *x* and *t*_0_, while *y* and *z* exhibited positive correlations. This pattern aligns with the spatial variations observed in other results, where *y* and *z* display similar trends, while *x* and *t*_0_ tend to increase from occipital towards central-frontal regions.

For these two features, the magnitude of the effect of *x* increased with posterior-anterior coordinates (e.g., moving from occipital to frontal regions), while the effect of *y* decreased, consistent with the predominance of corticothalamic activity in occipital regions. For alpha power, the magnitude of *z*’s effect decreased with spatial coordinates (coefficient Z:region = −9.256, *p <* 0.05), while the magnitude of *t*_0_’s effect increased (coefficient *t*_0_:region = 46.288, *p <* 0.05). Conversely, for the HF aperiodic exponent, the strength of the effect of *z* increased with space (coefficient Z:region = 1.574, *p <* 0.05) while *t*_0_’s decreased (coefficient *t*_0_:region = −32.295, *p <* 0.05). Regarding age, the magnitude of the effects of the parameters on alpha power remained similar except for *t*_0_, which increased with age (coefficient *t*_0_:age = 0.033, *p <* 0.05). For HF aperiodic exponent, all parameters had the magnitude of the effect increase with age except for *x* which did not significantly change with age (*p >* 0.05). MLM of alpha frequency showed a negative correlation with *x* and *t*_0_ strongest in occipital regions (coefficient *x* = −0.211, coefficient *t*_0_ = −6.002 *p <* 0.05). Parameters *y* and *z* have stronger effects in central-frontal regions which are, respectively, negatively and positively associated with frequency (coefficient *y* = −0.400, coefficient *z* = 0.442 *p <* 0.05).

By computing the cosine similarity of the brain saliences from both the behavioural PLS of the empirical features and modelling parameters, we can explore whether the regions identified by the empirical PLS are similar to the ones in the modelling PLS (Figure 5). Our analysis revealed that there is a high overlap between the regions identified by LV1 from the empirical PLS and LV2 from the modelling PLS. This suggests that the pattern of associations between frequency and ageing is similar to the associations between *t*_0_ and age in occipital regions and the limbic system. Conversely, regions identified by LV2 from empirical PLS overlapped with the ones identified by LV1 from modelling PLS, indicating a relationship between HF aperiodic exponent and *z* for central-frontal regions, including sensorimotor and prefrontal areas.

**Figure 5.**
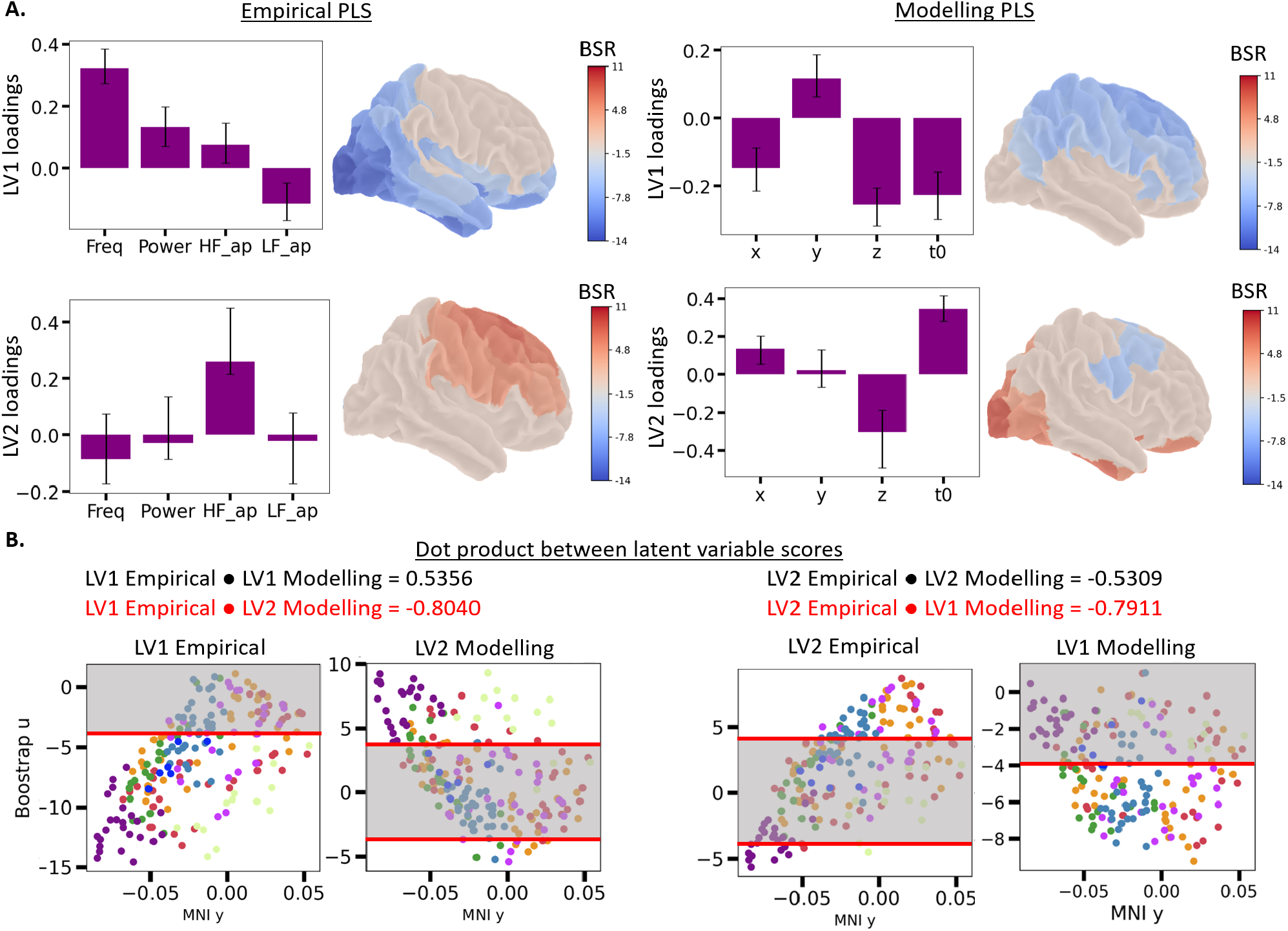
Comparison of empirical and modelling behavioural PLS results and the corresponding identified latent variables. **A)** Loadings of LV1 and LV2 from the empirical and modelling PLS, with the representation of the BSR on the brain. Empirical: LV1 presents high positive loadings of frequency, followed by power and is reliably showing a negative association with age for occipital regions and the limbic system. LV2 presents high positive loadings of HF aperiodic showing a reliable positive association with age in central-frontal regions. Modelling: LV1 has high loadings of z and *t*_0_ in the same direction, mostly showing negative relationship in central-frontal regions, meaning a positive association with z and *t*_0_. LV2 presents opposite directions for the loadings of z and *t*_0_ with a reliable positive association in occipital regions and limbic. **B)** Results of the cosine similarity between the brain scores u of the latent variables in both PLS, and the corresponding bootstrap ratios across the posterior-anterior axis. This suggest that the LV1 empirical is correlated with LV2 modelling and LV2 empirical is correlated with LV1 modelling.

Our results indicate that *z* and *t*_0_ are consistently varying with age across all features, with distinct spatial effects manifesting via alpha frequency, power and HF aperiodic exponent. These findings highlight the differential roles of corticothalamic and intrathalamic activities across different brain regions and their changes with age. This provides valuable insights into the neural mechanisms underlying resting-state oscillations.

## 1 Discussion

In this study, we investigated how resting-state power spectra vary across brain regions along the posterior–anterior axis and across the lifespan, linking these empirical patterns to underlying circuit mechanisms using a neurophysiological corticothalamic model. While previous studies have examined these aspects individually, our work is novel in integrating them within a unified framework. In particular, our modeling approach provides the first region-specific characterization of circuit dynamics underlying resting-state spectral variations with age as previous work explored it with a similar model in one channel [37]. We found distinct mechanisms varying across the cortex and age with higher corticocortical activity in frontal areas compared to posterior regions, and inversely for corticothalamic activity. We found that with ageing, posterior regions primarily exhibit increases in the corticothalamic delays and decreases in intrathalamic inhibition, while frontal regions present increases in the intrathalamic inhibitory circuitr, defining different spatial biomarkers.

### 4.1 Resting-state features present a posterior-anterior gradient with region-specific ageing effects

Our empirical results indicate that the alpha frequency gradient aligns with the trends reported in previous studies for similar age groups [53, 18, 26], showing a significant decrease along the posterior-anterior axis. However, we observe a loss of linearity in this gradient over ageing with a quadratic like relationship developing due to occipital regions experiencing a more pronounced decrease in frequency compared to central brain regions, despite a general decrease in frequency across most brain regions with age. This aligns with the results of Chiang et al. [32] which showed an alpha frequency decline particularly in occipital regions around 40-50 years of age. While alpha power, which demonstrates a strong correlation with posterior-anterior axis coordinate, declines slightly with age overall, reductions are more pronounced in occipital regions in line with previous reports [54, 55, 32]. As individuals age, their ability to process visual information and other sensory inputs declines [56, 57]. Caudal alpha rhythm has been suggested to be closely linked to sensory processing, including the regulation of visual and auditory information, and is also implicated in memory encoding and retrieval [58, 59, 60, 61]. Alpha activity is typically attenuated or blocked by increased attention and cognitive demands, particularly during visual tasks or other forms of mental effort [62, 63, 64]. Age-related reductions in alpha power may therefore reflect alterations in the neural mechanisms that support sensory information processing.

The HF aperiodic exponent is highest in posterior regions and decreases towards anterior regions, consistent with previous findings [21, 26, 22, 65]. Additionally, this exponent shows a spatial decline with age in occipital regions showing flatter slopes, as has also been observed in earlier studies during visual memory task [27] and resting (task-free) [65]. However, contrary to previous work [29, 65], we found certain frontal-central regions, including the dorsal and medial prefrontal cortex (e.g., PFCd and PFCmp subdivisions), the frontal medial cortex, with steeper slopes (increased HF aperiodic exponent) with age. Furthermore, we observed steeper slopes in the LF aperiodic exponent in posterior regions compared to frontal areas, contrasting with earlier reports that showed the opposite pattern [21, 22]. These differences could be explained by several factors. First, the methods and the frequency range used in these studies to estimate LF and HF differ (for example, LF exponent range either smaller between 0.1–2.5 Hz or wider 0.7-15 Hz), Our fitting of a monoexponentially decaying function encompasses a range up to 7Hz for the LF, while the HF was estimated using line fitting in the log scale in between 12-49Hz as they were not always well captured with FOOOF (Supplementary S.4). Furthermore, many previous analyses were conducted during eyes-open task-based conditions, focused on younger age ranges, and/or used different cortical parcellation schemes, which may further contribute to the observed discrepancies in aperiodic activity results [65].

Having characterized the empirical age-and region-related changes in oscillatory brain activity, we next sought to interpret these findings within a mechanistic framework. The main focus in the second half of the study was to offer a neurophysiological explanation for these empirically observed age and regional oscillatory brain activity changes.

### 4.2 Interpretation of corticothalamic and cortical contributions along the posterior–anterior axis

Our modelling analyses suggest that posterior regions are more driven by both corticothalamic (*y*) and intrathalamic (*z*) circuit activity than frontal areas, where corticocortical (*x*) loop gain is more prominent (Figure 3A). More specifically, an opposing spatial pattern is observed between corticothalamic and corticocortical circuit activity (Figure 3B), whereas intrathalamic circuit activity varies independently of these two. This underlying neurophysiological change is primarily observed in the power spectra as a decrease in alpha power, a change in the HF aperiodic exponent, and to a lesser extent the LF aperiodic exponent. A certain degree of anticorrelation between corticocortical (*x*) and corticothalamic loop gains (*y*) is expected, as both theory and experiments suggest that the brain operates near marginal stability, defined by the relation *x* + *y* = 1 [35, 49]. To maintain stability while supporting complex dynamics, the sum of *x* + *y* should therefore remain close to 1. Deviations far inside this boundary are thought to reflect states such as coma or deep anesthesia [33], whereas crossing the boundary is associated with transitions into large-amplitude seizure activity [66, 33]. Therefore, the minimal changes in *x* and *y* observed across ageing suggest that different brain regions remain within a consistent resting-state regime that is both stable and dynamically flexible, operating near-criticality [67]. This explains the opposite trend observed between the two loops. Increased corticothalamic circuit activity in posterior regions relative to anterior could reflect the stronger role of the thalamus in sensory areas. Corticothalamic loop activity in visual sensory areas is consistent with previous work that emphasized the role of the lateral geniculate nucleus (LGN) in the thalamus as a relay between the retinal fibers and the visual cortex [68, 69, 70]. Furthermore, prior work has suggested that sensory areas have more focal connections with the thalamus compared to associative areas which are more diffuse [71].

### 4.3 Regional changes in neural model parameters reflect age dependent mechanisms

With ageing, we observe increases in corticothalamic delay in occipital regions, coupled with a decrease in inhibitory intrathalamic loop gain. In contrast, fronto-central regions show increases in both intrathalamic loop gain and corticothalamic delay. These model parameter changes in ageing offer a candidate mechanistic explanation for empirical observations of the power spectra features. Specifically, frequency shows the strongest association with age, particularly in occipital regions, followed by the HF aperiodic exponent, which decreases in occipital regions and increases in fronto-central regions, as shown with our MLM and PLS results (Figure 4 and 5). It is worth noting, that among the studied circuit parameters, the intrathalamic gain was the only one to show opposing age-related trends across regions, namely between the visual and somatomotor network. This pattern suggests that distinct processes may underlie age-related changes in mu and occipital alpha rhythm.

Our findings are in alignment with Van Albada et al. [37], who investigated ageing variations by fitting a similar model on power spectra from the Cz electrode. They also noted an increase in the parameter *t*_0_ with age, which correlates with a reduction in the alpha frequency peak. This increase in corticothalamic delays could be attributed to the decline in thalamic white matter integrity with age [72]. The present study extends significantly the analysis of Van Albada et al. by examining closely the variation in spectral and estimated parameter features over the posterior-anterior axis. In our results, the age-related increase in *t*_0_ was most prominently observed in occipital regions. In the visual cortex, this was also accompanied by a decrease in HF aperiodic slope with reduced inhibitory intrathalamic circuit activity (*z*). This could be explained by pronounced atrophy with aging in posterior thalamic nuclei (LGN) [73], and/ or lower GABA levels in visual cortex compared to young adults [74].

Interestingly, we found increases in the *z* parameter in fronto-central regions with increased HF aperiodic exponent, indicating steeper slopes with ageing. In the last three deciles, (ages 58 to 88) the value of *z* becomes the highest in the somatomotor region. Similarly, Van Albada et al. [37] observed an increase in the absolute value of *G*_*srs*_ in the Cz electrode region, indicating heightened inhibitory intrathalamic loop gain. This is somewhat an unexpected result as with ageing thalamic volume declines [75, 76, 77] which would suggest overall lower activity. Several hypotheses could be proposed to explain the increased inhibitory intrathalamic circuit activity in those regions with ageing. First, some studies have shown that beta oscillations have been linked to the inhibitory neurotransmitter GABA which shows greater inhibitory activity within the motor cortices of older subjects [78, 79] but the opposite has also been suggested with reduced GABA levels [80, 81, 82, 83]. Second, a study reported an increased occurrence of beta events in older adults. Beta frequency (15–30Hz) rhythms are generated by inhibitory thalamocortical networks and are typically associated with an inhibited state of processing [84, 85]. In the context of our model, this pattern would be consistent with increased intrathalamic inhibition, suggesting that the somatosensory thalamocortical network in older adults may spend more time in a dampened state, potentially reducing its responsiveness and flexibility. Third, the increased intrathalamic inhibition could be explained by the increased expression of metabotropic glutamate receptors in the thalamus with ageing [86, 37]. Finally, studies have shown that the greatest volumetric shrinkage across the lifespan are observed in the caudate, cerebellum, hippocampus, and prefrontal regions, whereas the entorhinal the volume of the visual cortex presents minimal changes [87, 88]. A posterior-anterior gradient is also observed with age with greater structural deficiency in white matter located in frontal regions of the brain [89]. However, functional brain activity increases with age in frontal cortex which has been attributed to functional compensation [90, 88]. With our model, we observe increased intrathalamic inhibitory activity which could represent a compensatory mechanism whereby the thalamic reticular nucleus enhances inhibition of relay neurons to filter increasingly noisy frontal cortical feedback. While the precise mechanisms remain unclear, the model captured distinct ageing-related changes between occipital and fronto-central regions, most notably differentiating occipital alpha and central mu rhythms, with *z* capturing these neurophysiological variations. Furthermore, this highlights that therapeutic brain stimulation targeting strategies may need to differ depending on the region of interest, as the underlying healthy mechanisms vary. Our results also raise the possibility of using *z* as a region-specific biomarker for tracking the effects of both disease progression and treatment interventions.

### 4.4 Limitations

It is important to recognize some limitations of this study. Firstly, despite having a substantial sample size encompassing a wide range of ages, a longitudinal approach would better facilitate tracking individual patterns of spontaneous neural activity throughout the lifespan. While our cross-sectional design enabled the examination of a broad age spectrum, it did not allow for the assessment of individual-specific developmental trajectories. With regards to the model, although our neural field model offers a comprehensive overview, it may oversimplify the complex nature of neurobiology, providing a general understanding without fully capturing the detailed interactions within the brain. This simplification is advantageous for modelling, as it allows us to focus on the essential complexities necessary to explain observed phenomena. However, it inherently limits the model’s ability to represent the full intricacies of brain function.

### 4.5 Conclusion

In conclusion, our study elucidates significant spatial variations in resting-state in alpha power and aperiodic components, revealing a pronounced gradient which may be attributed to the distinct roles of corticothalamic and corticocortical activities in posterior and frontal regions, respectively. We identified that ageing is associated with a decline in alpha frequency, power, and the HF aperiodic exponent in occipital regions, likely due to increased corticothalamic delay. In contrast, fronto-central regions show increased HF aperiodic exponent, suggesting elevated inhibitory intrathalamic circuit activity. The modelling results reveal spatial differences related to age, indicating distinct mechanisms of neural ageing across various brain regions. Our findings enhance the understanding of age-related neural dynamics and pave the way for improved predictive models and targeted neurological treatments based on M/EEG resting-state measurements. Future research should focus on longitudinal studies to track individual neural activity patterns over time to glean further insights on the complex interactions underlying these age-related changes. Additionally, the association between cognitive scores and modelling results could be investigated to determine which features across the brain serve as stronger biomarkers.

## Data and code availability

The analysis and visualization code used in this paper is included in this GitHub Repository https://github.com/GriffithsLab/Bastiaens2025_CorticothalamicRestingParametrization

The maxfiltered MEG data used in this work is from the Cam-CAN project which are available from http://camcan-archive.mrc-cbu.cam.ac.uk, subject to conditions specified on the website. For a complete description of Cam-CAN data and pipelines, please refer to Shafto et al. (2014) and Taylor et al. (2017).

The MCMC model fitting algorithm implemented on MATLAB is available at https://github.com/BrainDynamicsUSYD/braintrak.

## Author Contributions

S.B.: Conceptualization, Formal analysis, Methodology, Visualization, Writing – original draft, Writing – review & editing, D.M: Conceptualization, Methodology, Writing – review & editing, L.R.: Methodology, Software, Writing – review & editing, T.M: Conceptualization, Software, Writing – review & editing, K.K.: Conceptualization, Writing – review & editing, M.P.O.: Conceptualization, Writing – review & editing, J.D.G: Conceptualization, Funding acquisition, Methodology, Supervision, Writing – review & editing.

