## Supplementary information for "Corticothalamic circuit mechanisms underlying brain region and ageing variations in resting-state alpha activity"

### S.1 Results of full analytical model parameters including PC, PCA and PLS

As a complement to the main text, this section includes the full analysis of the gain parameters from the analytical model, as well as $\alpha$ and $\beta$, which were omitted from the main figures for clarity and interpretability.

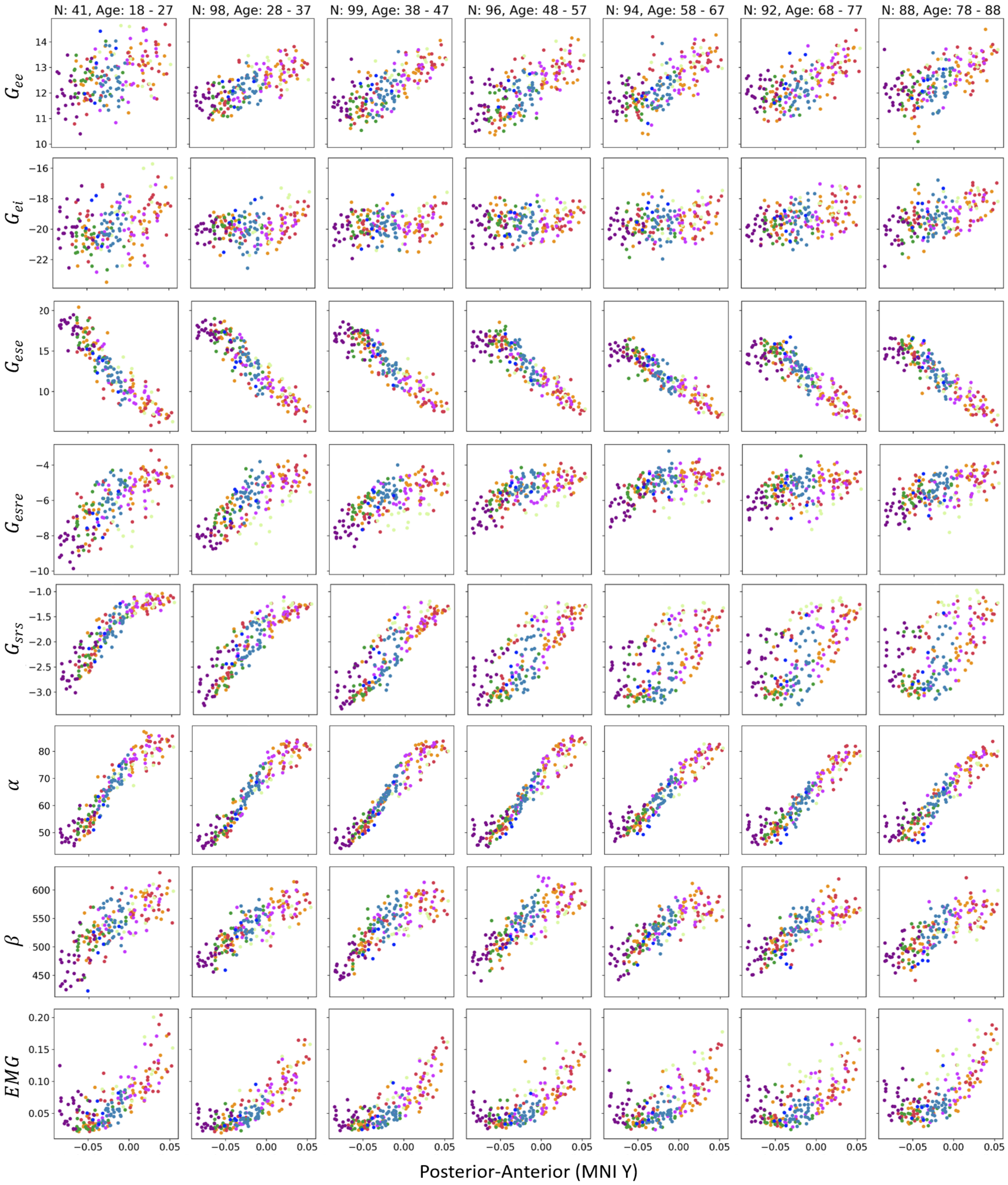


**Figure S.1A.** ***Spatial gradient of full analytical model parameters along the posterior-anterior axis with ageing variations*** The two parameters that most consistently show spatial trends across age groups are $G_{ese}$ and $\alpha$, following patterns similar to those observed in alpha power. In occipital regions, ageing is associated with a marked reduction in inhibition, as reflected by decreases in $G_{esre}$. Finally, we observe a spreading of the gain $G_{srs}$ with age, following a similar pattern to HF aperiodic activity.

Table S.1. Summary of Empirical Population-level Pearson Correlation Results

| Age Range | $G_{ee}$ | $G_{ei}$ | $G_{ese}$ | $G_{esre}$ | $G_{srs}$ | $\alpha$ | $\beta$ | $EMG$ |
|---|---|---|---|---|---|---|---|---|
| 18 - 27 | r = 0.4445 | r = 0.3322 | r = -0.9221 | r = 0.7281 | r = 0.9068 | r = 0.9420 | r = 0.7696 | r = 0.7308 |
| 28 - 37 | r = 0.7081 | r = 0.2599 | r = -0.9250 | r = 0.7711 | r = 0.8915 | r = 0.9562 | r = 0.8397 | r = 0.8206 |
| 38 - 47 | r = 0.7462 | r = 0.1976 | r = -0.9169 | r = 0.6539 | r = 0.8807 | r = 0.9599 | r = 0.8154 | r = 0.8050 |
| 48 - 57 | r = 0.7073 | r = 0.1515 | r = -0.9028 | r = 0.6473 | r = 0.7940 | r = 0.9577 | r = 0.8309 | r = 0.7453 |
| 58 - 67 | r = 0.7031 | r = 0.2819 | r = -0.9203 | r = 0.5642 | r = 0.6233 | r = 0.9495 | r = 0.8263 | r = 0.6492 |
| 68 - 77 | r = 0.6356 | r = 0.4183 | r = -0.8798 | r = 0.3783 | r = 0.6073 | r = 0.9378 | r = 0.8046 | r = 0.6133 |
| 78 - 88 | r = 0.6201 | r = 0.5319 | r = -0.9090 | r = 0.5819 | r = 0.5788 | r = 0.9336 | r = 0.7919 | r = 0.6860 |
| p-value | p<<0.001 | p<<0.001 | p<<0.001 | p<<0.001 | p<<0.001 | p<<0.001 | p<<0.001 | p<<0.001 |
| Percentage of significant correlations | 21.38% | 14.14% | 72.53% | 38.98% | 81.58% | 95.07% | 49.84% | 72.53% |

The spatial distribution of the full gain parameters reveals significant spatial correlations for most subjects, particularly for $\alpha$, $G_{srs}$, $G_{ese}$, and $EMG$ (with 95.07%, 81.58%, 72.53%, and 72.53% of subjects respectively). The most consistent spatial correlations across all age ranges are seen with $G_{ese}$ and $\alpha$, both showing r-values greater than 0.8. In contrast, the spatial correlation for $G_{srs}$ diminishes progressively with age, as r-values decline from 0.9068 in the youngest group to 0.5788 in the oldest, consistent with the age-related changes in the $z$ parameter. Even though a spatial correlation with $x$ has been previously identified, only a small number of subject present a significant correlation in the posterior-anterior axis for $G_{ee}$ and $G_{ei}$, which could be explained by the high variability of $G_{ei}$ across subjects as said in vanAlbada. Details of all the r-values, p-values and the percentage subject with a significant correlation is summarized in Table S.1.

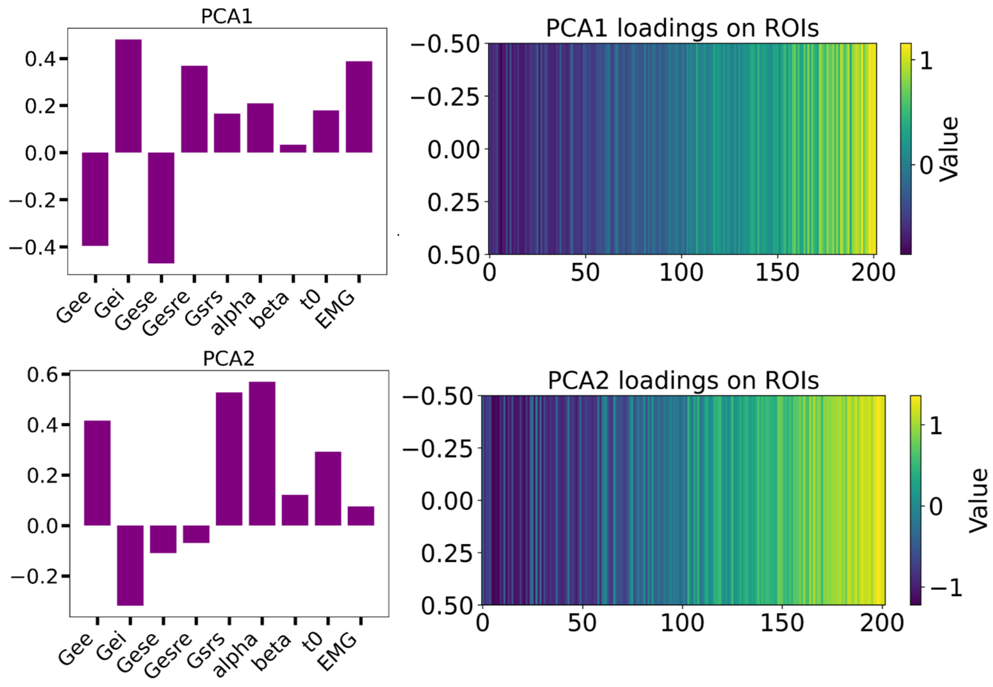


**Figure S.1B.** ***PCA results*** The first PCA component is negatively loaded with $G_{ee}$ and $G_{ese}$, and positively loaded with $G_{ei}$ and $G_{esre}$. However, the last two values are negative which means that they are more inhibitory in the opposite trend.

The first two components of the PCA analysis with the parameters of the analytical model with all the gains explain 54.5% of the variance. In the first component (PCA1), we observe that corticothalamic-related parameters (such as $G_{ese}$ and $G_{esre}$) have a stronger influence in occipital regions, comparable to the spatial distribution of the $y$ parameter (Figure S.1B). In the second component, $G_{srs}$ and $\alpha$ show strong positive loadings. Spatially, $G_{srs}$ is more negative in occipital regions, while $\alpha$ is more positively loaded in frontal regions. Additionally, the opposing loadings of $G_{ee}$ and $G_{ei}$ in both PCA components suggest an inverse relationship between these parameters: when $G_{ee}$ has a more positive loading, $G_{ei}$ tends to have a more negative loading, and vice versa. Since $G_{ei}$ is a negative value, this indicates that both $G_{ee}$ and $G_{ei}$ exhibit high magnitudes concurrently, but in opposite directions. This may indicate a balance between excitation and inhibition within the corticothalamic circuit.

| Feature | $G_{ee}$ | $G_{ei}$ | $G_{ese}$ | $G_{esre}$ | $G_{srs}$ | $\alpha$ | $\beta$ |
|---|---|---|---|---|---|---|---|
| Corr. with LV | 0.5130 | 0.4313 | 0.1928 | 0.2463 | 0.3918 | 0.2679 | 0.4065 |

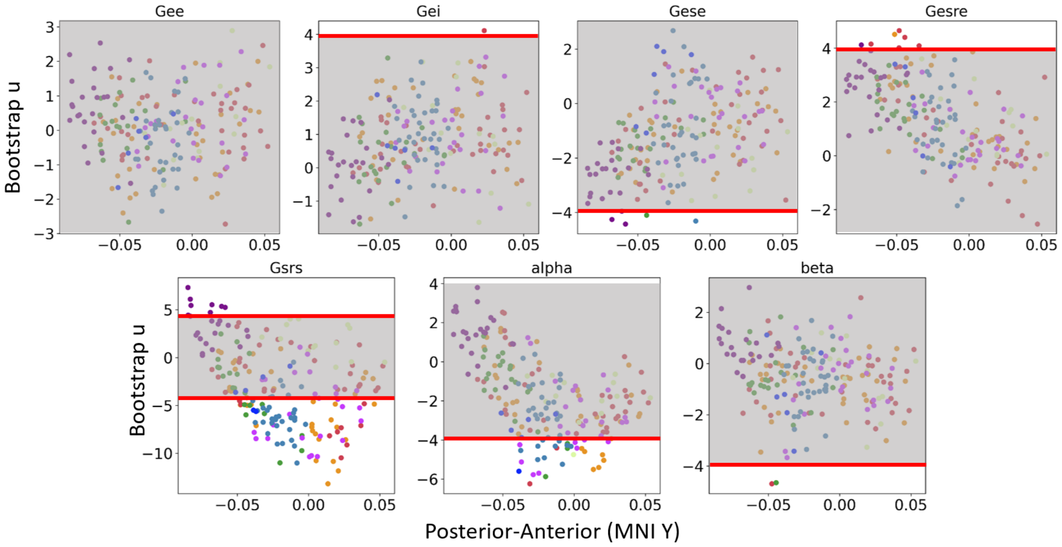


**Figure S.1C.** ***Behavioural PLS results*** Bootstrap ratios for each region over the posterior-anterior axis for full model parameters. Changes with age are reliably seen in most regions in parameters $G_{srs}$, and $\alpha$.

The PLS analyses identified a significant relationship with age for all parameters except for $G_{ee}$ (with $p = 0.0999$, above the significance threshold of $p = 0.05$). For all other parameters, $p << 0.05$. However, this relationship is considered reliable in most regions only for $G_{srs}$ and $\alpha$, based on their bootstrap ratios. This indicates that most regions age-related changes from $z$ are due to $G_{srs}$ changes, followed by $\alpha$ in central-frontal regions, in addition to the ageing changes presented with $t_0$ in main text.

#### S.2 Details and full output of MLM analyses

We provide in details the results from the mixed linar model analyses that are summarized in the Figure 4 and section 3.6.

Table S.2A. Alpha Frequency Mixed Linear Model Regression Results

| | |
|---|---|
| Model: MixedLM | Dependent Variable: Alpha Frequency |
| No. Observations: 121400 | Method: REML |
| No. Groups: 607 | Scale: 0.4290 |
| Min. group size: 200 | Log-Likelihood: -122422.7041 |
| Max. group size: 200 | Converged: Yes |
| Mean group size: 200.0 | |

| | Coef. | Std.Err. | z | $P > \|z\|$ | [0.025 | 0.975] |
|---|---|---|---|---|---|---|
| Intercept | 10.839 | 0.095 | 113.634 | 0.000 | 10.652 | 11.026 |
| x | -0.215 | 0.071 | -3.034 | 0.002 | -0.353 | -0.076 |
| age | -0.007 | 0.002 | -3.923 | 0.000 | -0.010 | -0.003 |
| x:age | -0.016 | 0.001 | -13.244 | 0.000 | -0.018 | -0.013 |
| y | -0.400 | 0.080 | -4.980 | 0.000 | -0.557 | -0.242 |
| y:age | 0.001 | 0.001 | 0.419 | 0.675 | -0.002 | 0.003 |
| z | 0.448 | 0.089 | 5.010 | 0.000 | 0.273 | 0.623 |
| z:age | 0.011 | 0.001 | 7.686 | 0.000 | 0.008 | 0.014 |
| t0 | -6.046 | 0.497 | -12.160 | 0.000 | -7.021 | -5.072 |
| t0:age | 0.058 | 0.009 | 6.699 | 0.000 | 0.041 | 0.075 |
| region | 26.015 | 0.551 | 47.174 | 0.000 | 24.934 | 27.096 |
| x:region | -33.955 | 0.559 | -60.766 | 0.000 | -35.050 | -32.859 |
| y:region | 4.976 | 0.664 | 7.489 | 0.000 | 3.674 | 6.278 |
| z:region | 8.543 | 0.628 | 13.594 | 0.000 | 7.311 | 9.775 |
| t0:region | -111.921 | 4.132 | -27.084 | 0.000 | -120.020 | -103.821 |
| Group Var | 0.300 | 0.027 | | | | |

Table S.2B. Alpha Power Mixed Linear Model Regression Results

| | |
|---|---|
| Model: MixedLM | Dependent Variable: Alpha Power |
| No. Observations: 121400 | Method: REML |
| No. Groups: 607 | Scale: 0.0273 |
| Min. group size: 200 | Log-Likelihood: 44351.3297 |
| Max. group size: 200 | Converged: Yes |
| Mean group size: 200.0 | |

| | Coef. | Std.Err. | z | $P > \|z\|$ | [0.025 | 0.975] |
|---|---|---|---|---|---|---|
| Intercept | 1.090 | 0.035 | 30.774 | 0.000 | 1.020 | 1.159 |
| x | -0.423 | 0.018 | -23.691 | 0.000 | -0.458 | -0.388 |
| age | -0.004 | 0.001 | -6.766 | 0.000 | -0.005 | -0.003 |
| x:age | 0.002 | 0.000 | 6.023 | 0.000 | 0.001 | 0.002 |
| y | 0.492 | 0.020 | 24.265 | 0.000 | 0.452 | 0.532 |
| y:age | 0.001 | 0.000 | 2.155 | 0.031 | 0.000 | 0.001 |
| z | 0.801 | 0.023 | 35.415 | 0.000 | 0.756 | 0.845 |
| z:age | -0.008 | 0.000 | -22.409 | 0.000 | -0.009 | -0.007 |
| t0 | -4.042 | 0.126 | -32.175 | 0.000 | -4.288 | -3.796 |
| t0:age | 0.033 | 0.002 | 15.343 | 0.000 | 0.029 | 0.038 |
| region | -10.629 | 0.139 | -76.353 | 0.000 | -10.901 | -10.356 |
| x:region | 5.360 | 0.141 | 37.999 | 0.000 | 5.083 | 5.636 |
| y:region | -6.032 | 0.168 | -35.966 | 0.000 | -6.361 | -5.703 |
| z:region | -9.256 | 0.159 | -58.342 | 0.000 | -9.567 | -8.945 |
| t0:region | 46.288 | 1.043 | 44.377 | 0.000 | 44.244 | 48.333 |
| Group Var | 0.060 | 0.021 | | | | |

Table S.2C. LF Aperiodic Mixed Linear Model Regression Results

| | |
|---|---|
| Model: MixedLM | Dependent Variable: LF Aperiodic |
| No. Observations: 121400 | Method: REML |
| No. Groups: 607 | Scale: 0.1875 |
| Min. group size: 200 | Log-Likelihood: -72197.5537 |
| Max. group size: 200 | Converged: Yes |
| Mean group size: 200.0 | |

| | Coef. | Std.Err. | z | $P > \|z\|$ | [0.025 | 0.975] |
|---|---|---|---|---|---|---|
| Intercept | 0.639 | 0.063 | 10.081 | 0.000 | 0.514 | 0.763 |
| x | -0.132 | 0.047 | -2.818 | 0.005 | -0.223 | -0.040 |
| age | 0.004 | 0.001 | 4.026 | 0.000 | 0.002 | 0.007 |
| x:age | -0.007 | 0.001 | -9.519 | 0.000 | -0.009 | -0.006 |
| y | -0.013 | 0.053 | -0.237 | 0.813 | -0.117 | 0.091 |
| y:age | -0.003 | 0.001 | -3.672 | 0.000 | -0.005 | -0.002 |
| z | -0.123 | 0.059 | -2.087 | 0.037 | -0.239 | -0.007 |
| z:age | 0.016 | 0.001 | 16.374 | 0.000 | 0.014 | 0.017 |
| t0 | 3.244 | 0.329 | 9.867 | 0.000 | 2.599 | 3.888 |
| t0:age | -0.012 | 0.006 | -2.167 | 0.030 | -0.024 | -0.001 |
| region | -1.734 | 0.365 | -4.774 | 0.000 | -2.455 | -1.026 |
| x:region | 16.831 | 0.369 | 45.509 | 0.000 | 16.089 | 17.537 |
| y:region | -4.874 | 0.439 | -11.094 | 0.000 | -5.735 | -4.013 |
| z:region | -11.839 | 0.415 | -28.496 | 0.000 | -12.653 | -11.025 |
| t0:region | -91.234 | 2.732 | -33.395 | 0.000 | -96.588 | -85.879 |
| Group Var | 0.133 | 0.018 | | | | |

Table S.2D. HF Aperiodic Mixed Linear Model Regression Results

| | |
|---|---|
| Model: MixedLM | Dependent Variable: HF Aperiodic |
| No. Observations: 121400 | Method: REML |
| No. Groups: 607 | Scale: 0.1084 |
| Min. group size: 200 | Log-Likelihood: -39127.8238 |
| Max. group size: 200 | Converged: Yes |
| Mean group size: 200.0 | |

| | Coef. | Std.Err. | z | $P > \|z\|$ | [0.025 | 0.975] |
|---|---|---|---|---|---|---|
| Intercept | 2.606 | 0.060 | 43.792 | 0.000 | 2.490 | 2.723 |
| x | -1.453 | 0.036 | -40.854 | 0.000 | -1.523 | -1.383 |
| age | -0.004 | 0.001 | -3.485 | 0.000 | -0.006 | -0.002 |
| x:age | 0.001 | 0.001 | 1.442 | 0.149 | -0.000 | 0.002 |
| y | 0.703 | 0.040 | 17.431 | 0.000 | 0.624 | 0.783 |
| y:age | -0.003 | 0.001 | -4.731 | 0.000 | -0.005 | -0.002 |
| z | 2.321 | 0.045 | 51.583 | 0.000 | 2.233 | 2.409 |
| z:age | 0.003 | 0.001 | 4.472 | 0.000 | 0.002 | 0.005 |
| t0 | -2.088 | 0.250 | -8.349 | 0.000 | -2.578 | -1.598 |
| t0:age | 0.033 | 0.004 | 7.591 | 0.000 | 0.024 | 0.041 |
| region | -2.237 | 0.277 | -8.069 | 0.000 | -2.780 | -1.694 |
| x:region | 0.226 | 0.281 | 0.804 | 0.421 | -0.325 | 0.776 |
| y:region | -2.803 | 0.334 | -8.391 | 0.000 | -3.457 | -2.148 |
| z:region | 1.574 | 0.316 | 4.984 | 0.000 | 0.955 | 2.194 |
| t0:region | -32.295 | 2.077 | -15.548 | 0.000 | -36.366 | -28.224 |
| Group Var | 0.152 | 0.027 | | | | |

#### S.3 Motivation for frequency-specific aperiodic fitting of LF and HF slope

In this supplementary section, we present the rationale for independently estimating LF and HF slope instead of using the estimated FOOOF fitting results.
Figure S.1 shows a selection of power spectra fitted using our custom LF and HF slope estimation method, followed by comparison with FOOOF's default 'fixed' and 'knee' aperiodic fitting modes. These examples illustrate that the 'fixed' FOOOF mode adequately captures the LF slope but fails to fit the slope above approximately 12Hz, and is still affected by it, which skews the LF slope estimation. In contrast, the 'knee' mode better accounts for higher frequencies, yet does not capture the LF slope. Furthermore, the 'knee' fitting was not successful across all PSDs. It is worth noting that our HF line fitting can be influenced by large spectral peaks in the beta band; however, this approach provided the best compromise between flexibility and robustness across subjects.

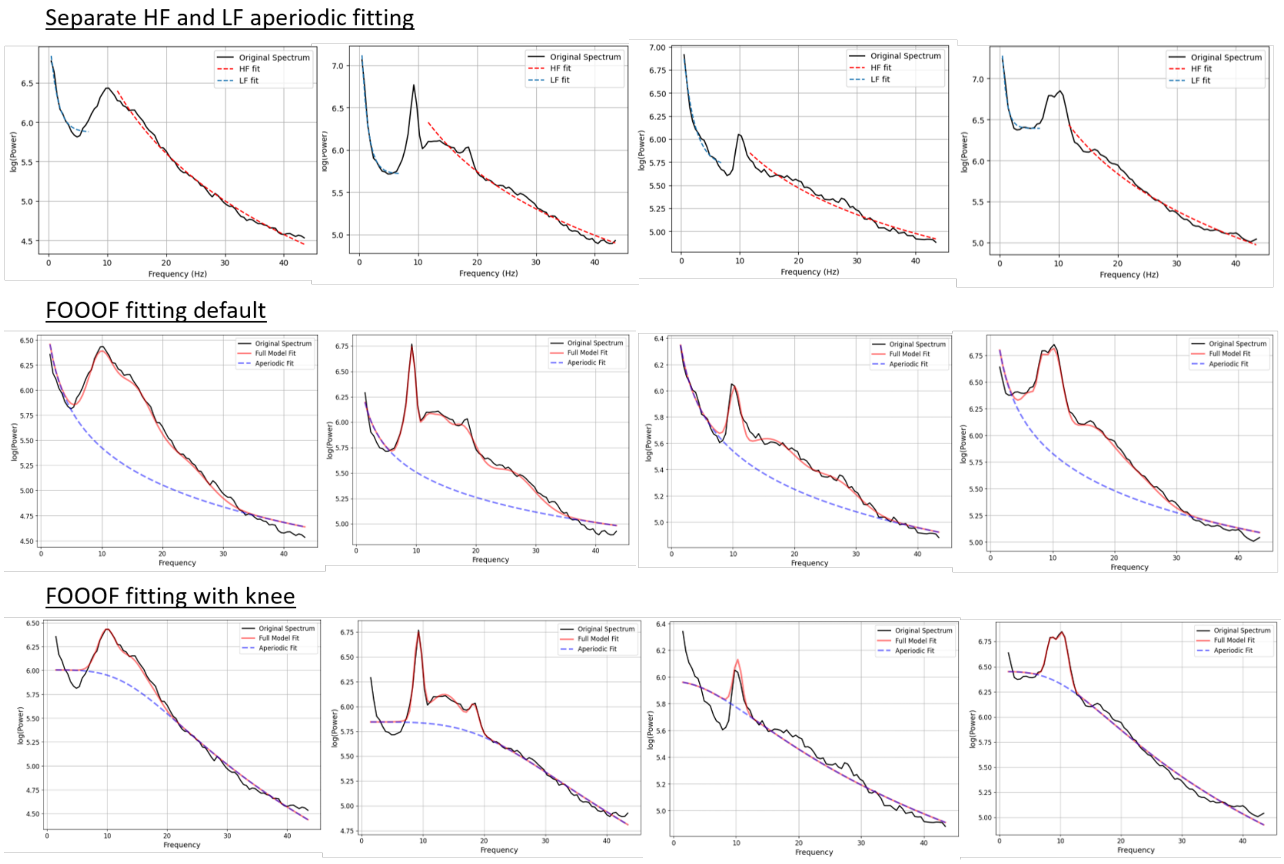


**Figure S.3A.** ***Select PSD fits using FOOOF across different subjects.*** PSDs from four representative subjects are shown, each fit using our custom method, and then FOOOF with both the default (no-knee) and knee-enabled settings. The default method primarily aligns with the LF slope, while the knee-enabled method tends to better capture the HF components. This highlights a tradeoff in fitting accuracy across the frequency spectrum, depending on the FOOOF configuration used with our dataset, and supports the rationale for estimating separate slopes for low and high frequencies rather than a single unified slope.

We still present here in the following section the results over the posterior-anterior axis for different age groups using the exponents estimated with FOOOF over space and age to assess the differences with our own analysis. Compared to the 'knee' fitting, we observe similar trends in the main analysis for HF aperiodic activity except for the somatomotor regions which present steeper slopees with age. As mentioned this difference could be due to the fact that our method is skewed by high beta peaks which might be more prevalent in this region. For the default FOOOF analysis, which is affected by LF and HF slope, we see an overall slight decrease in exponent is observed till 48 years old which then seems to increase even in frontal regions resulting in slightly positive correlations (that are not significant). For most age ranges, the spatial gradient is not significant (Table S.3A)

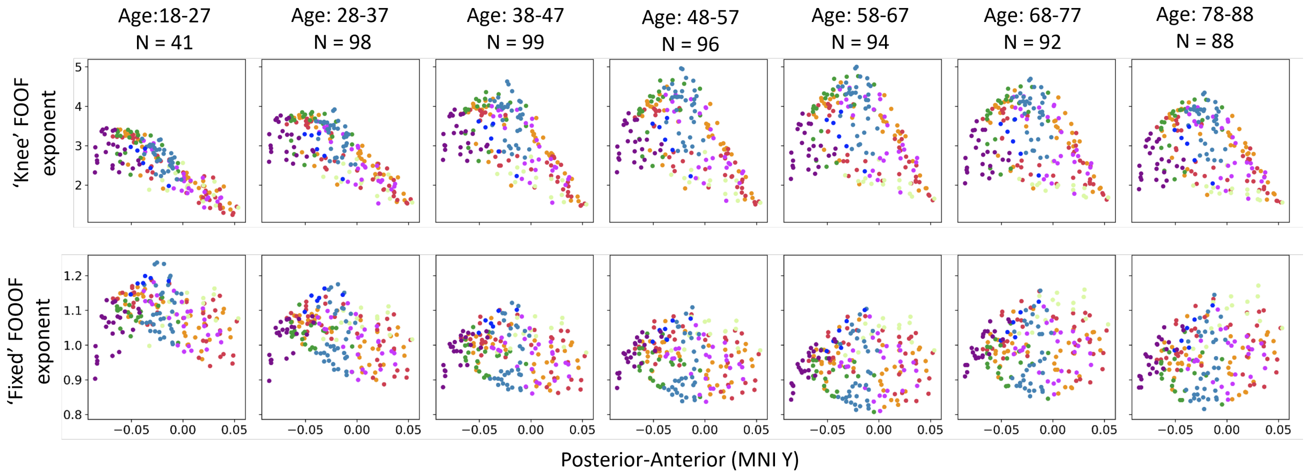


**Figure S.3B.** ***Spatial gradient of exponent estimated with FOOOF methods***. The upper plot corresponds to the FOOOF exponent estimation using the 'knee' method which follows more the slope of the higher frequencies (above 12Hz). The plot below is the exponent using the default parameter options ('fixed') which tends to follow the LF.

Table S.3A. Summary of Empirical Population-level Pearson Correlation Results

| Age Range | Aperiodic HF | Aperiodic LF |
|---|---|---|
| 18 - 27 | r = -0.838 | r = -0.212 |
| 28 - 37 | r = -0.758 | r = -0.283 |
| 38 - 47 | r = -0.648 | r = -0.179 |
| 48 - 57 | r = -0.513 | r = -0.107 |
| 58 - 67 | r = -0.397 | r = 0.041 |
| 68 - 77 | r = -0.385 | r = 0.108 |
| 78 - 88 | r = -0.412 | r = 0.212 |
| p-value | p<<0.001 | p>0.001 (NS, except for range 28-37) |
